# The hedonic evaluation of neurofeedback stimuli is fast, automatic and implicit An ERP study on stimulus design

**DOI:** 10.64898/2026.06.06.730592

**Authors:** Adrian Naas, Danpeng Cai, Payam S. Shabestari, Tobias Kleinjung, Delphine Ribes Lemay, Patrick Neff, Andreas Sonderegger

## Abstract

Introduction - Neurofeedback (NFB) has demonstrated efficacy in treating various disorders, often achieving substantial symptom reductions. Despite its effectiveness, a significant percentage of users (non-responders), fail to benefit from NFB. Addressing this issue, the study at hand investigates the role of NFB design on neuro-physiological responses. Method - event related potentials (ERPs) are examined in response to the application of different aesthetic principles in the context of NFB stimulus design. Drawing from Self-Determination Theory and ERP studies in the field of web design, 16 feedback stimuli were developed according to specific design principles. Stimulus design effects were inspected by means of ERPs allowing for the assessment of implicit and fast electroencephalogram (EEG) reactions. Results of n = 38 participants indicated distinct ERP response patterns at time window of interest 1 (TWOI-1; 100-200 ms) and TWOI-2 (200-300 ms), predicted by beholder-based liking and complexity of stimuli. The findings align with the proposed hypotheses suggesting that aesthetic evaluation of NFB stimuli occurs rapidly and implicitly. In response to the aesthetic vs. non-aesthetic categories, the findings were mixed. The results underscore the importance of further exploration of aesthetic design guidelines in the context of NFB applications. It is discussed how NFB aesthetics relate to Processing Fluency and Affective Prediction Error Theory, while the theoretical methodological issue of the Fixed Effect Fallacy is taken into consideration. The study contributes to the broader understanding of how design elements can affect therapeutic efficacy and engagement in NFB and Human Computer Interaction in therapeutic contexts in general.

## Introduction

EEG Neurofeedback-Training shows important effects on various disorders like Attention Deficit Hyperactive Disorder [1,2] and Tinnitus [3]. However, current NFB procedures struggle with non-responder rates ranging between 16% and 57% [4] – users failing to learn neuromodulation as a result of their NFB practice [5]. The number of users profiting from the treatment could potentially be increased by addressing NFB performance, via aesthetic pathways [6]. At the moment, it is not understood how NFB stimulus aesthetics relate to neural activity patterns (potential NFB neural targets) and to mental health outcomes. It is the subject of this study to evaluate whether the effect of NFB stimuli aesthetics on ERP correlates is fast, automatic and implicit. The current lack of research in NFB stimulus design can be attributed to the complexity of the NFB closed-loop (see Fig 1). Evaluating directional effects of one closed loop element another (e.g. visual feedback effects on the user) in real time (Lubianiker et al., 2019) is methodologically taxing [6].

**Fig 1.**
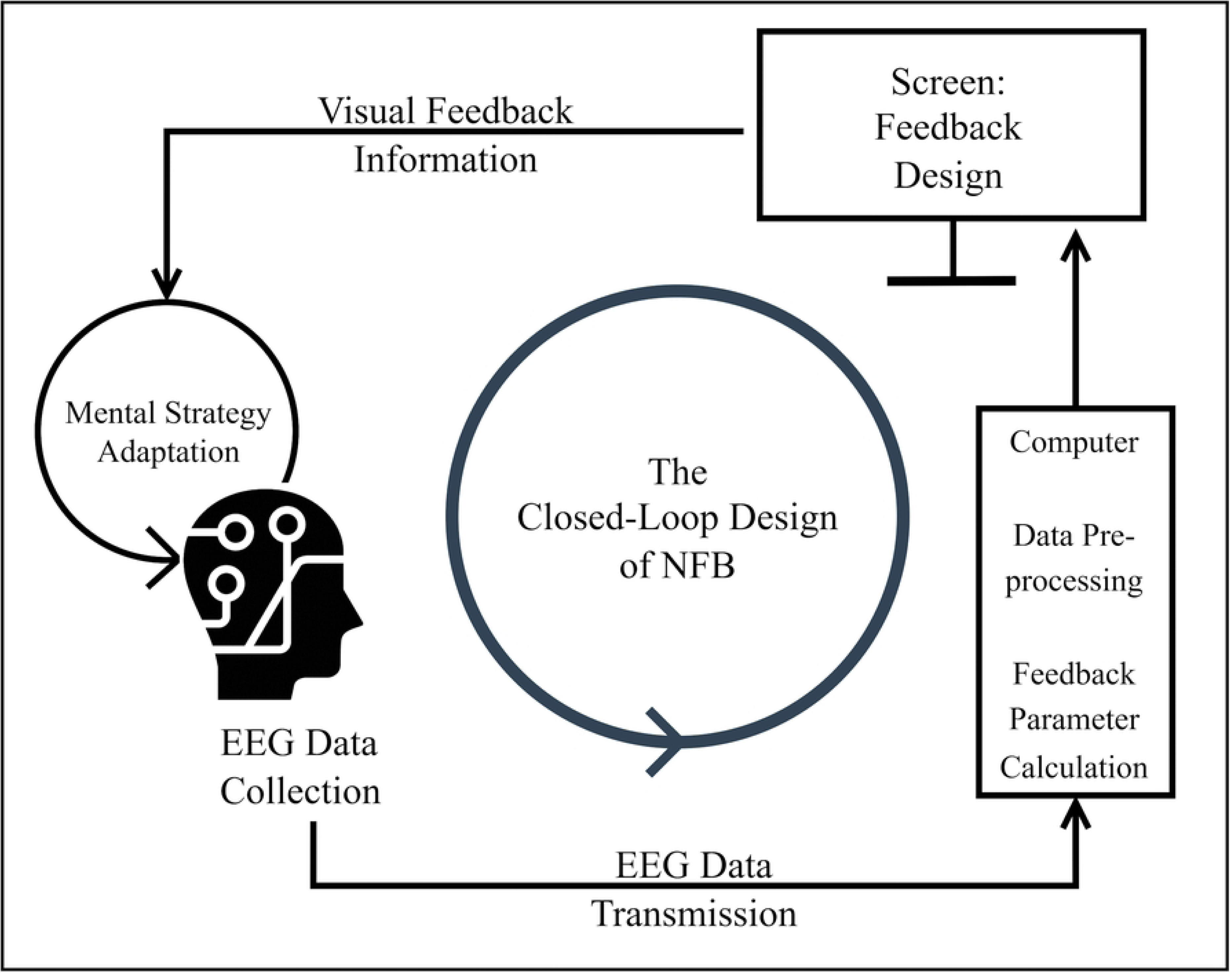
The Closed Loop Design of Neurofeedback Training. Abbreviations in the graph are electroencephalogram (EEG) and neurofeedback training (NFB).

In order to assess possible effects, it is crucial to isolate closed loop elements experimentally by applying useful research designs. E.g. ERP studies allow for the quantification of EEG responses per stimulus [7]. Such studies are essential as stimulus design might be a covariate of NFB training success - i.e. the target brain activity pattern might be influenced by the stimulus itself.

There is e.g. ERP evidence [7] that samples of individually disliked logos and liked logos are processed differently. More specifically, differences in the ERP response were found in an early TWOI between 150-200 ms (central-parietal P1) and at TWOI 200-400 ms at central and parietal locations (central N2 and parietal-occipital P2). Handy et al. [7] concluded that the individual hedonic evaluation of mundane stimuli like logos is fast (takes place before 200 ms) and implicit (without conscious aesthetic judgement). Note, the observed effects of aesthetics in logo design might be due to stimuli being designed to provoke fast automatic pleasure responses [7].

Note, hedonic evaluations have to be differentiated from aesthetic experiences [8]. However, in the field of Neuroaesthetics, these two processes have been shown to be strongly interlinked [9]. At the moment, distinct neural processes or regions of interest (ROIs) for hedonic appreciation of a stimulus (i.e. liking) and aesthetic experiences cannot be identified [9]. In the same vain, Chatterjee and Vartanian [10] argue that aesthetic experiences are related to a state of disinterest interest – identified by liking without wanting. That is why liking is considered an indicator of aesthetic experience in the article at hand.

Research [11–13] similar to Handy et al. (2010) seems to confirm respective effects taking place especially 100-200 after stimulus onset (TWOI-1) and clear evidence can be found for TWOI-2 between 200-300 ms. Notably, the directionality of observed ERP effects is, to date, somewhat inconclusive. For example, in response to pictures of every-day objects, Righi et al. [12] found smaller P2 amplitudes for liked stimuli in the fronto-parietal network within Go Stimuli in a Go-Nogo Task. Note, P2 was defined between 150-250 ms in anterior locations and in posterior locations between 190-290 ms. Simultaneously, the authors found larger P2 amplitudes for NoGo stimuli in the same time range. Additionally, liked stimuli elicited larger P1 (50-150 ms) components at tempo-occipital locations.

In a study on ERP effects of Chinese typeface aesthetics, Li et al. [11] found no differences in ERPs with respect to the P1 or N1 (150-200 ms) component, however, an effect of the P2/N2 component (200-300 ms) of liking was apparent. The effect was based on ERP grand average differences and disliked Chinese typefaces seemed to produce more pronounced P2/N2 components – an effect that is discussed by the authors in terms of the line of argument of Herbert et al. [14] indicating that more pronounced P2 components are commonly observed in stimuli being perceived as very positive or very negative.

Adding to the current literature, Wang et al. [13] conducted a study in the context of marketing layout interfaces. No effects of preference were found for the P1/N1 (150-200 ms) component but, again, with respect to the P2 component (200-300 ms), a main effect of preference was found pointing at differences between disliked and other stimuli. Summing up, there seems to be some evidence for liking resulting in altered ERP trajectories between 100-200 ms (TWOI-1) relating to P1 and N1 components. The evidence is clearer with respect to effects taking place between 200-300 ms (TWOI-2) and, thus, relating to the P2 and N2 components.

As results are still somewhat inconsistent with regard to early component effects [for a review, see 15] it is crucial to explore respective effects in the context of NFB stimuli. The lack of current efforts to evaluate established routes of performance increase is surprising given the empirical evidence in the domain of human computer interaction studying effects and consequences of design aesthetics. In this regard, various studies have shown beneficial effects of aesthetics on individuals’ motivation, perseverance and performance [6,see e.g., 16–20].

With regard to performance, aesthetics effects seem to be especially relevant in situations of high task difficulty [21,22] – a crucial finding in the context of NFB as non-responders receive little positive feedback. More specifically, the mechanism results from the NFB principle of contingent reward in response to performance. Arguably, NFB users with low entry motivation and the concurrent lack of finding a successful mental NFB strategy is mediated by a lack of positive reinforcement in NFB users with low performance levels. The vicious circle of amotivation creating non-responders in the process [6] might be buffered by aesthetics effects on perseverant behaviours and performance [6,16,19,20,23].

In the light of these effects, it seems reasonable to assume that design aesthetics might play an important role in NFB. Given the potentially important influence of design aesthetics in NFB, the research gap of non-existent NFB design guidelines is surprising [6,24], underlining the importance of studying consequences of design aesthetics in NFB stimuli.

While the presented effects of aesthetic experiences on ERP trajectories [7,11–13,15] helps to describe a noteworthy relationship - it does not explain the underlying mechanisms of the effect. As an explanatory framework Mere Exposure Theory can be consulted. It is well established that aesthetic experience interacts with situations of repeated stimuli encounters [25]. With increasing exposures to a stimulus, aesthetic experiences are improved as a function of the number of exposures [26].

The Mere Exposure Effect is supported by a robust body of literature and the empirical evidence has been reviewed recently by Montoya et al. [25]. The authors point to the fact that the relationship between encounters and aesthetic experience is not linear and follows an inverted U-shape trajectory.

I.e. with increasing numbers of encounters boredom is increased and the aesthetic experience diminishes [25]. The inverted U trajectory of exposure and aesthetic experiences is further modified by the complexity of a given stimulus. Simplicity leads to a faster and steeper slope of increase (and decrease) of exposures on aesthetic experiences, while complexity results in a gentler up-wards slope and downwards slope.

Apart from complexity interactions with aesthetic design, repeated encounters and liking, it has been shown that the concreteness of a stimulus is connected to stimulus aesthetics [27]. Concreteness can be defined as the level to which a visual stimulus is the representation of people or real-world materials or objects [28]. In abstract visualisations, however, meaning is transported with shapes, lines and symbols like arrows. Thus, the link to real world objects is less clear in abstract representations [29].

While aesthetics effect dynamics are well established in other contexts, there is a scarcity of studies on NFB stimulus aesthetics effects [6,24]. That is why it seems critical to establish a general research procedure to approach the endpoint of aesthetics and design effects on NFB learnability (Reiner et al., 2018) and symptom outcomes [e.g. Tinnitus suffering;, 3].

A useful approach (especially in quantitative research in comparison to qualitative research) is to begin investigating a new area by conducting studies in a simplified context, where the set of prerequisite conditions is still relatively broad (see Fig 2). This allows fundamental processes to be identified before narrowing the focus. For example, before claiming that the aesthetics of visual NFB designs influence NFB learning and symptom outcomes, one must first establish a more basic prerequisite: That visual design in general can produce measurable effects on neural responses.

**Fig 2.**
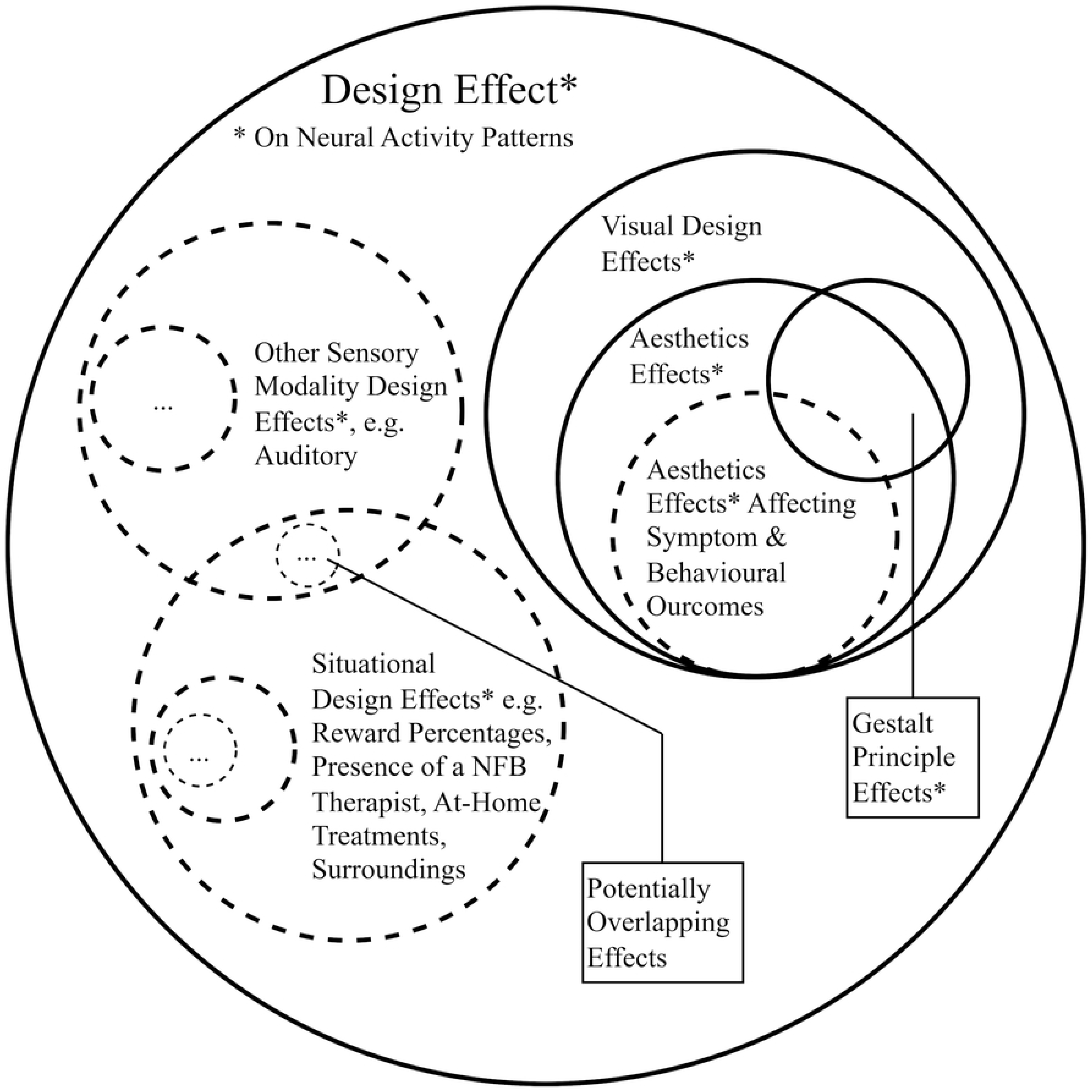
Venn Diagram of Potential Design Effects Sorted According to Specificity. Continuous lines displaying target effects of the article at hand. Abbreviations in the graph are electroencephalogram (EEG) and neurofeedback training (NFB).

Laying the groundwork for studies on aesthetics effects on NFB performance and, thus, potential effects mitigating the non-responder issue, the research question is formulated: Does aesthetics in NFB stimuli produce fast, automatic and implicit ERP correlates? As aesthetics is linked to stimulus complexity in the situations of repeated exposure [25], we hypothesise that ERPs vary as a function of complexity and liking, possibly following the inverted u-shape trajectory described by Montoya et al. [25].

In order to adopt controlled experimental procedures [27], different design dimensions were manipulated leading to stimuli comprising a large variance in design aesthetics, complexity and concreteness. It was the secondary aim of this study to test whether aesthetic design principles affect neurophysiological outcomes and participant ratings similar to beholder-based aesthetics i.e. liking ratings.

## Materials and method

### Participants

Advertising on campus and mailing lists were used as participant recruitment strategies. Recruitment started on the 7^th^ of December 2023 and it ended on the 9^th^ of October 2024. After being informed about study contents and the possibility to terminate participation at any point without the obligation to provide a reason, all participants provided their written informed consent. The final number of participants was planned to be comparable to the study of Handy et al. [7] who used a sample size of *N* = 32. The final sample consisted of 38 participants (27 female, 11 male). Participants ranged in age between 18 and 33 years (*M* = 22.58 years, *SD* = 3.94 years).

The study was conducted in three languages, 14 participants completed the study in English, 10 participants followed study procedures in French and 14 participants took part in German. The participants indicated not being diagnosed with any psychiatric or neurological disorders - a statement that resonated with low Beck depression inventory fast screen values, indicating low levels of psychological distress in the sample (sum score *M* = 2.42, *SD* = 3.59) with a mean value similar to measurements reported in the general population [30]. All participants indicated normal or corrected to normal vision. None of the participants had any prior experience with NFB. Psychology students were compensated by means of points on a university intern credit system. Non-psychology student participants received a bar of chocolate. Ethical approval was granted from the internal review board of the University of Fribourg/Freiburg (reference No. 2022-802 R1).

### Experimental protocol

The recruitment of participants took place mainly via announcements on the official experiments platform at the University of Fribourg as well as via advertising of the study during psychology lectures for first-year students. Further participants were recruited among the acquaintances of study personnel. Finally, study flyers were distributed around the university campus.

The inclusion and exclusion criteria for participation were informed consent, fluent language level of one of the study languages (German, English or French), no existing neurological and psychiatric disorders, a minimum age of 18 years, normal or corrected-to-normal vision and no previous experience with NFB. The exclusion of participants with previous NFB experience ensured that aesthetic judgements of NFB stimuli independent from possible familiarity effects. The random allocation of the test subjects to the respective conditions took place using a randomised block design (e.g. for the control variable "up" or "down" in the context of the distraction task, see Fig 3).

**Fig 3.**
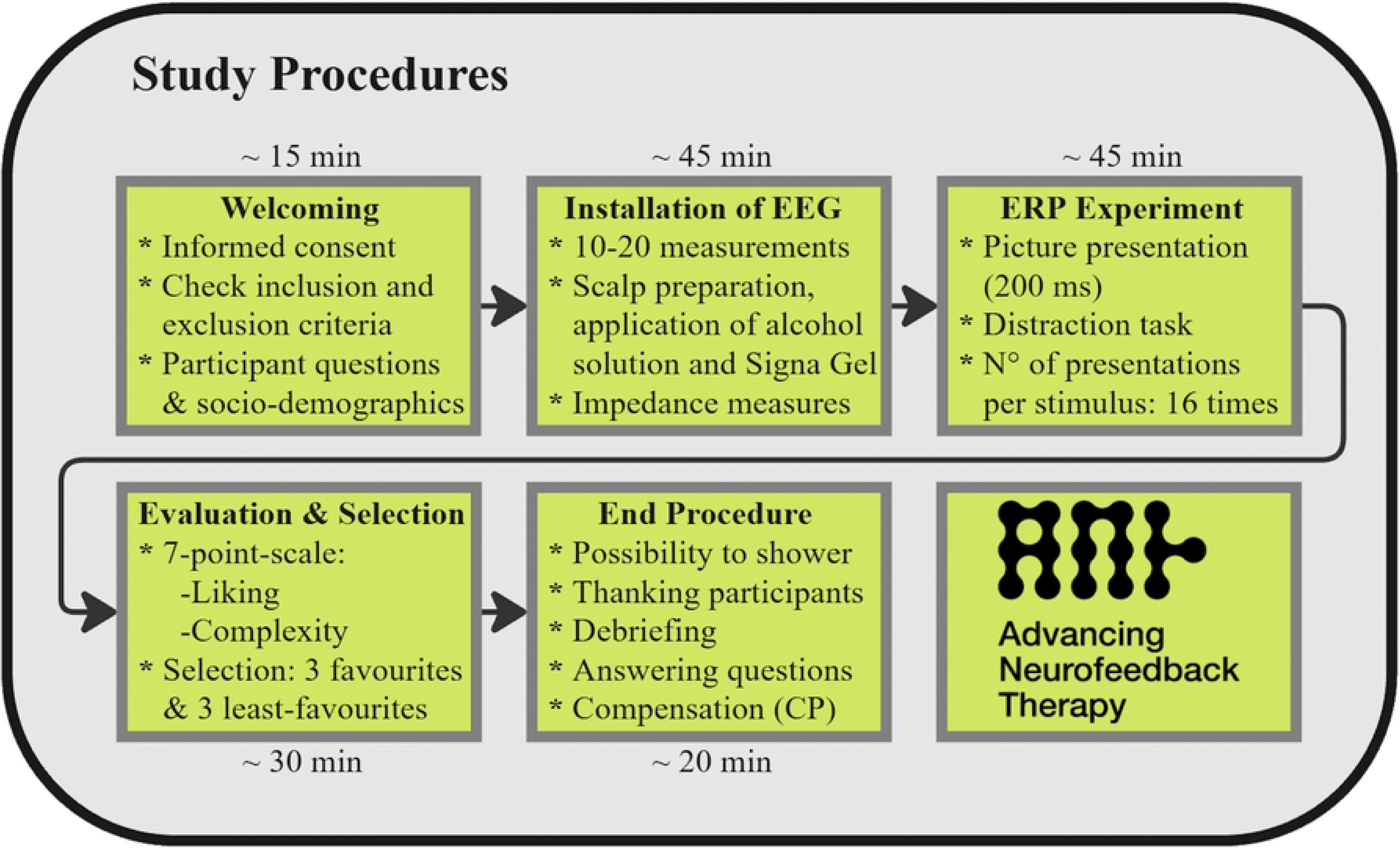
Study Procedures in the Laboratory. Abbreviations in the graph are electroencephalogram (EEG), event-related potential (ERP) and credit points (CP).

Overall, the experiment took about 2 hours and 35 minutes to complete. It comprised an introduction and informed consent, the installation of the EEG (see section *Materials and Equipment*), the ERP task (see section *The ERP Task*), different scales for the evaluation of NFB prototypes and an end procedure allowing participants to shower, follow the debriefing and receive the compensation. During debriefing, the aim of the distraction task was explained to the participants and the study aim was described: The evaluation of implicit automatic EEG reactions to different design stimuli.

### Measures and instruments

Liking scores were collected per NFB prototype and assessed by means of a seven-point-scale (1 = “Dislike”, 7 = “Like”), similar to Handy et al. (2010). Analogously, we adopted the approach suggested by Handy et al. [7] for the assessment of complexity and the applied scale ranged from 1 (= “simple”) to 7 (= “complex”). Finally, Preference was assessed by asking participants to indicate which picture was their first, second or third favourite (e.g. “Which picture was your favourite? Please choose 1 picture. You make a selection by clicking on the respective picture”). Least-Preference was measured similarly.

### Equipment

Throughout the experiment, data collection was facilitated by the use of two computers. Computer 1, a Dell Latitude 7420, Windows 10 Enterprise Version 22H2 was connected to the participants screen (Samsung Odyssey, 62.2cm, Nvidia G-Sync, resolution 1920 x 1080 pixels, 240 Hz refresh rate) and was employed to present the ERP study paradigm as well as questionnaires. The connection between the high-frequency screen and the Dell Latitude 7420 was created with a high-speed fibre optic HDMI 2.0 cable. The mouse and keyboard were also connected to the computer via high-speed cables. A second computer of the same type and Windows version ran the EEG recording software (OpenBCI graphical user interface; GUI) recorded the data with a sampling rate of 125 Hz as text files (.txt) that were stored on a local hard drive for subsequent data analyses. Questionnaires were administered via the online survey software Unipark, version EFS 21.2.

EEG signals were registered using a low-impedance electrode gel (ECI Electrode Gel) and a small (50-54cm), medium (54-58cm) or large (58-62cm) 21-channel OpenBCI EEG cap. The caps comprise proprietary, sintered Ag/AgCL electrodes and the cap was connected with Touch-Proof connectors (1.5 mm) to the 16-channel OpenBCI Cyton + Daisy PCBs (16 channels + reference and ground). The OpenBCI Cyton, as well as the Daisy board involve 8 high gain / low noise input channels [31], have a 24-bit data resolution per channel and are connected passively to the cap. PCBs incorporated the Texas Instruments ADS1299 ADC. Voltage input of the boards can vary between ∼ 3.3 - 12 V and Rechargeable batteries were applied (Panasonic NCR18650B, 3350 mAh) and recharged between test sessions with the Lipo Rider Plus, Charger/Booster - 5V/2. It is an advantage of EEG operation on batteries that the technology reduces electrical line noise (frequencies at 50 Hz). Instead of using the Cyton board’s built-in micro SD-card slot, the data was sent to the Bluetooth dongle via the RFduino Low Power Bluetooth radio. The dongle comprises the FT231X USB-to-serial converter from FTDI and a RFD22301 radio module by RFdigital.

Electrodes sites were chosen according to international 10-20 placement system: FP1, FP2, AFz, F3, F4, F7, F8, Cz, C3, C4, T3, T4, Pz, P3, P4, O1, O2. The averaged activity over M1 and M2 was set to be the reference and ground was assigned to AFz. Note, the reference electrodes were attached independently from the cap with separate electrodes of the same material (Ag/AgCL). To guarantee low resistance EEG contact quality electrode positions were prepared with alcohol-soaked cotton swabs. Additionally, conductive gel (eci ELECTRO-GEL) was applied. For the duration of the experiment impedances were maintained at levels ≤ 15 kOhm. Overall participants, 3 electrodes were interpolated due to impedances 15>kOhm.

### EEG preprocessing and data analysis

Recorded raw EEG data underwent preprocessing using a custom-built Matlab 2022a pipeline [32], incorporating EEGLAB [33] and ERPLAB [34] plugins. The pipeline included EEGLAB default FIR bandpass filter, *pop_eegfiltnew()*, with cut-off frequencies between 1 and 30 Hz. This procedure removed power-line noise and as a result, no additional notch filter at the 50 Hz was applied.

Visual inspections of power density plots and raw EEG showed the procedure confirmed the absence of line noise. In terms of filtering, a hamming window was applied while the filter order was set to default (automatic) - i.e. transition bandwidth and filter order were derived by means of a heuristic. More precisely in the context of applied bandpass filters, transition bandwidth was defined as 25% of the lower passband edge frequency and a minimum of 2 Hz to ensure stability. In some circumstances, this criterion cannot be met - e.g. due to the proximity of the passband to a specific frequency (e.g. Nyquist frequency, defined as half the sampling rate or 0 Hz). For these cases, the transition bandwidth was defined as distance between bandpass edge and that specific frequency. The approach was applied to warrant feasible filter design and minimising edge effects as well as spectral distortion.

In order to exclude artifacts of eye movements, an independent component analysis (ICA) approach was used. EEGLAB [33] offers by default an ICA decomposition and subsequent component classification as artifact or brain activity. The classification was taken into account during the visual inspection and components of eye movements and blinking were rejected respectively.

Then, the data were sectioned into non-overlapping ERP 1100 ms epochs with baseline set between -100 and 0 seconds relative to stimulus presentation. The following artifact rejection was carried out with *eeg_regepochs()* removing artifact-prone epochs automatically via *pop_autorej()*. The function applies iterative standard deviation-based thresholding identifying epochs with improbable or extreme amplitudes. ERPs were calculated as mean amplitudes over TWOI-1 and TWOI-2.

Epoching according to the trigger input and the marker .txt file was carried out in order to identify respective trials in the EEG data (e.g. prototype 1). Then, epochs were sorted into the categories of interest (e.g. category “non-aesthetic prototypes”, see Fig 4) and the ERP curves were calculated and plotted as the mean of all trials over all participants in that category utilising the *std_erpplot()* function [33]. Plots were generated using a custom Matlab script (see Supplementary Materials A) including a 30 Hz filter. Finally, for the multivariate pattern analysis [MVPA; 35] predicting stimulus category based on ERP activity, the MNE suite [36] and MNE-Python [37].

**Fig 4.**
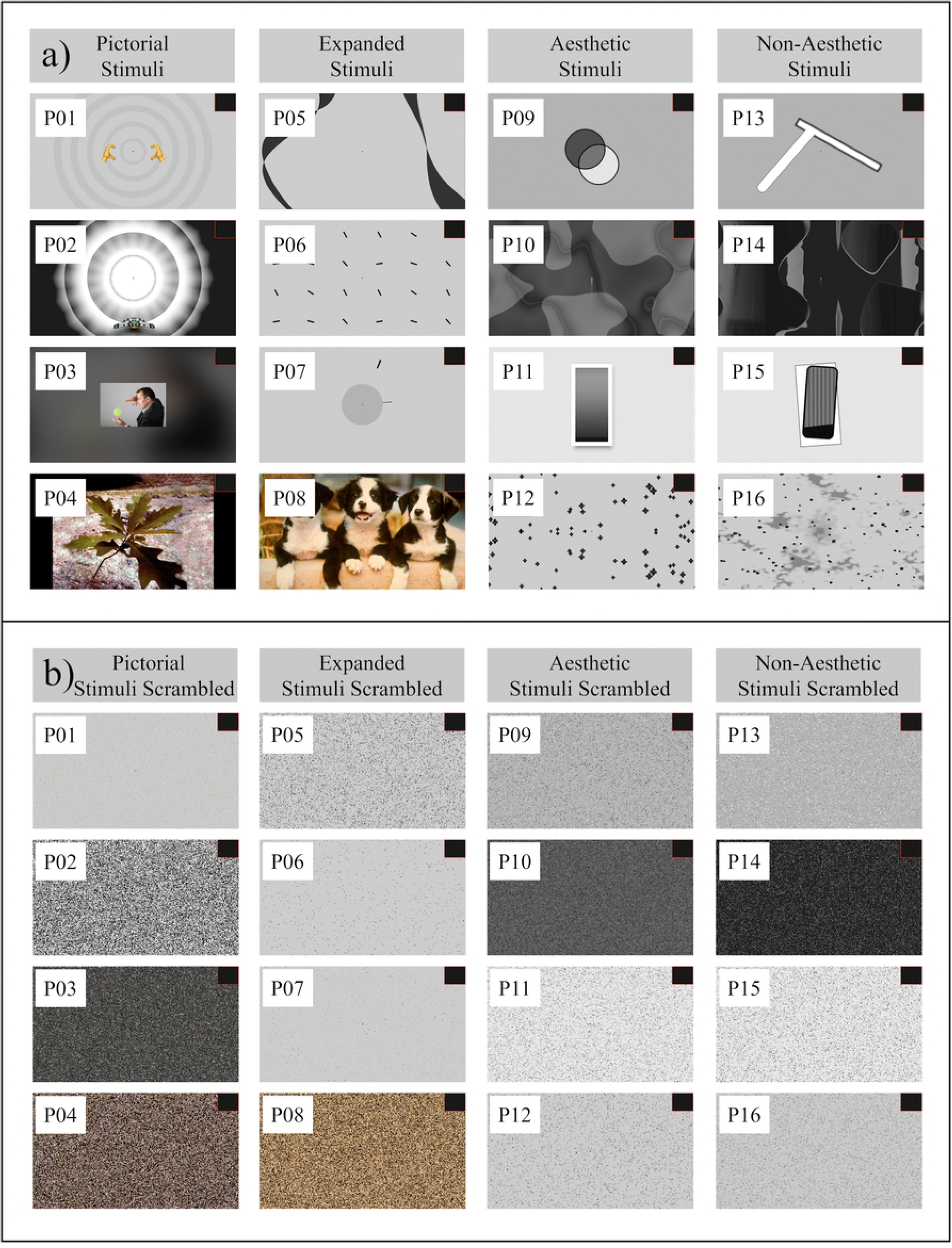
ERP-Stimuli. Stimuli are depicted in the Categories “Concrete”, “Expanded”, “Aesthetic” and “Non-Aesthetic” as well as their scrambled counterparts. Abbreviations in the graph are event-related potential (ERP).

In order to verify the time output measures generated by the software Presentation [38] a physical flicker procedure was applied in the study at hand. Decreases in illumination on the screen were captured by means of a photosensor and had the effect of increasing the respective resistance. Electrical resistance was monitored constantly with a frequency of about 10 kHz by an Arduino UNO R3 microcontroller and a voltage of 200 µV (the standard output of 5.0 V was attenuated by means of a voltage divider circuit) was sent to the analogue inputs of the Cyton+Daisy board.

Thus, the presentation of a stimulus was indicated by a concurrent voltage jump from 0 to 200 µV in the analog read of the OpenBCI GUI recording. At the same time Neurobehavioural Systems Presentation wrote stimulus presentation times into a log file. The two systems were synchronised during offline analyses by lining-up the first increase in voltage of the analog read in EEG recording with the presentation time of the first stimulus of the Neurobehavioural Systems Presentation log file. The specific EEGLAB functions for those procedures were *pop_chanevent()* to create markers according to the increases in voltage on the analogue read input of the OpenBCI Cyton board text file and *pop_importevent()* to import and align the log file event data.

### The ERP task

To ensure the assessment of implicit and fast hedonic evaluation of NFB stimuli, participants followed a distraction task which consisted of a simple reaction time task. Participants were advised to make a fast decision on whether the presented stimulus (200 ms presentation time) was a NFB prototype or a scrambled image. Decision results were indicated by clicks of arrow keys on the keyboard. More specifically, the presentation of a NFB prototype was indicated by clicking the “Up” arrow key and the presentation of a scrambled NFB prototype was communicated by clicking the “Down” arrow key. Note, the key assignment (e.g. scrambled stimulus → Up) was counterbalanced.

Pictures of the 16 prototypes were presented 30 times per participant. The same presentation frequency per participant was applied for scrambled stimuli. A fixation point (colour hexa decimal code #05648C) was shown for the whole duration of the ERP experiment. Background colour was set to white and study instructions were presented with black font colour. Stimuli were optimised in terms of central position in order to keep ERP lateralisation to a minimum.

### Stimuli

Four categories were established, representing varying degrees of aesthetic appeal and concreteness. These categories were used to transport the message of “you’re doing good” in the context of NFB - when participants achieve the NFB goal (e.g. successful increase of alpha amplitudes in alpha NFB) they need to be notified and receive a respective operant conditioning reward. According to communication theory [39] such messages can be transmitted in different ways and via different mediums and in NFB mediums are diverse ranging from computer games and meditation-like applications to simple thermometer representations. To capture a wide range of possible mediums, 16 NFB prototypes were created that can be divided into the four categories (1) aesthetic, (2) non-aesthetic, (3) concrete and (4) expanded prototypes, henceforth called *manipulated prototype categories*.

To create the aesthetically pleasing stimuli, principles drawn from “Gestalt Psychology” [40] were applied. The word "Gestalt" translates to form, pattern, or shape in German. The theoretical framework of “Gestalt” Psychology emphasises the notion that perception is based on organised patterns and that we see whole images rather than collections of parts. According to previous empirical work on icon design [27], the application of Gestalt theory comprises a focus on principles like symmetry, similarity and “Prägnanz” [40]. Non-aesthetic designs were created by inversing Gestalt principles and applying asymmetry, dissimilarity and missing distinction between background and foreground.

The third stimulus category consisted of concrete designs incorporating figurative and photorealistic content. As all other prototypes were abstract and as concreteness has been shown to affect aesthetics appeal and liking [27], the manipulated prototype category “concrete” was added. In that sense the prototype set mirrors the current NFB design landscape comprising, at times, heavily gamified and concrete stimuli. For example, the thermometer prototype concreteness was increased by showing the picture of a magician levitating a ball.

As it is the secondary aim of the study at hand to compare theory-guided NFB prototypes to the current standard in NFB prototype design. That is why the designers explored gut feelings and design hunches - giving rise to an extension of the stimulus set by four prototypes in the manipulated category “expanded”. Note, designers were given the possibility to discuss with NFB and EEG specialists beforehand yielding the following design guidelines: Central focus to reduce eye movement, evolving fluidic forms to maintaining prolonged visual interest, communicating positive reinforcement via synchronisation to induce satisfaction, emphasising feelings of “satisfying rewards”. Finally, the possibility to use selected affective picture of the International Affective Picture System [IAPS; 41] comprising high levels of valence, and provoking positive emotions intrinsically was proposed. A variety of four prototypes within each level of the manipulated prototype categories was developed (see Fig 5).

**Fig 5.**
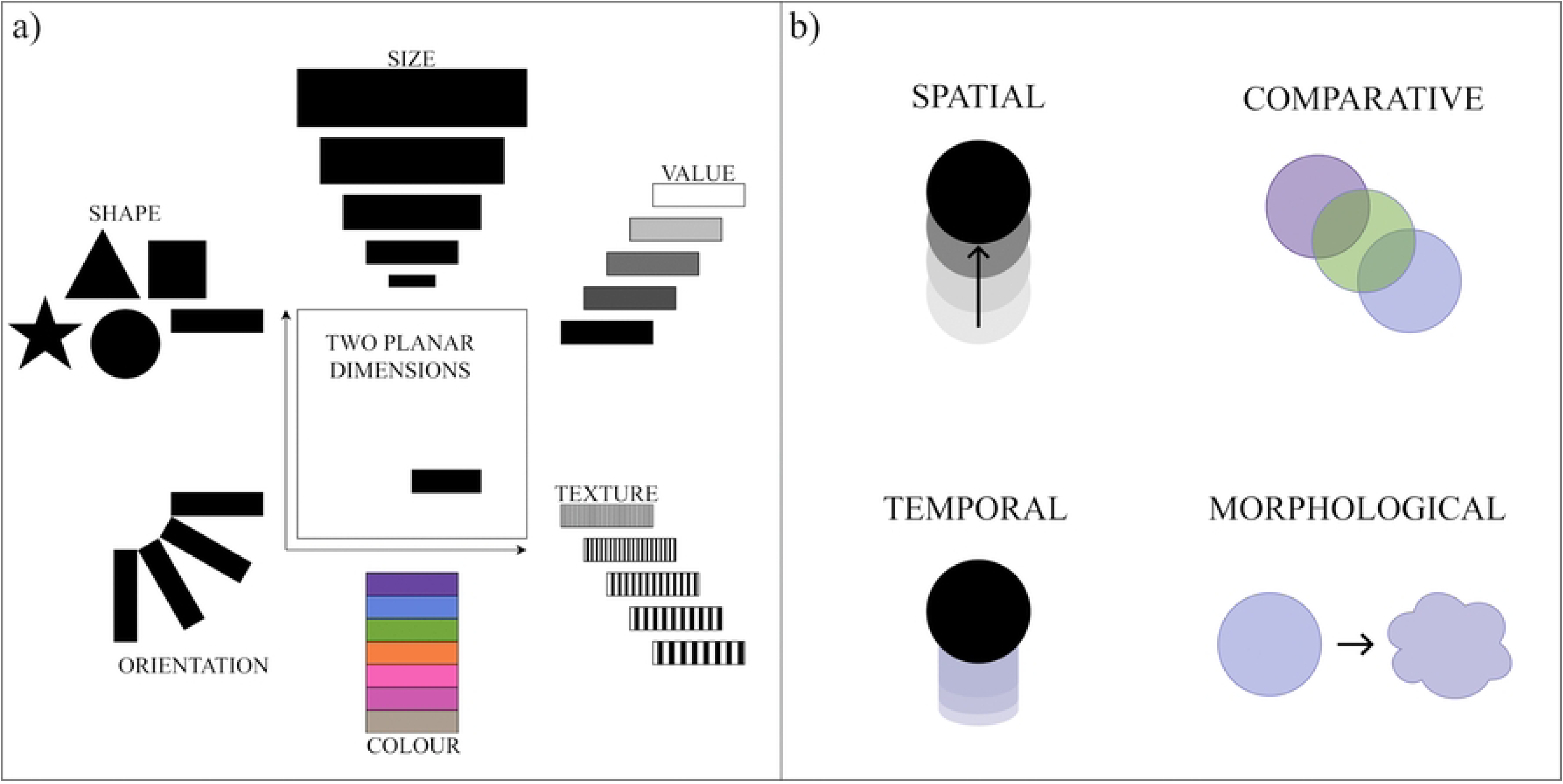
Stimulus Design Concepts. Visual feedback categories (panel a) and their scrambled counterparts (panel b), drawing inspiration from the work of Jacques Bertin (1983).

More specifically, within each manipulated prototype category, one prototype drew inspiration from thermometer designs commonly used in NFB [see e.g., 42,43]. Thermometer prototypes used attributes such as size, volume or location to convey the NFB message (e.g. “producing higher alpha amplitudes” leads to a higher level of the thermometer bar). A second variation was created by applying a comparative or leveller approach. It involved comparing multiple visual elements and overlapping shapes. E.g. the desired brain could be indicated by increasing the overlap of two dynamically moving spheres (see prototype 9). In the third variation, the concept of object accelerating over time over time and/or the achievement of a screen position was explored. A final, fourth variation was created by involving transitions of one geometrical shape or picture into another.

### Pretests

In order to streamline experimental procedures and to train student personnel, 28 participants were tested in advance. In the process, fixation cross colour was changed in order to make the experiment more comfortable to the eyes of the participants, the visual synchronisation system with a flicker channel and photo sensor was improved. Visual stimuli were adapted in terms of central positioning. Furthermore, the distraction task was adapted in order to require participants to actively process the stimuli. The final task constituted a simplified procedure of the Simon task [44] requiring participants to react to properties of the stimuli themselves i.e. the stimulus being a “design” or a “scramble” (see Fig. 4, panel b). Other adaptations of procedures in the pretest process concerned different translations of the study instructions (translation - back translation procedures were applied), as well as reference electrode placement (changed from earlobes to mastoids).

### Statistical analysis

Statistical Analyses were performed using r-studio version 4.5.1 [45]. A general procedure was selected modelling the hypothesized effects in the context of a multilevel linear model with the r-package “lme4” [46] and R [47]. The models allowed for random intercepts, as including the term allows participants to vary within conditions in terms of y-axis intercepts. E.g. assuming an effect of liking ratings on ERP amplitudes, participants vary randomly in terms of their ERP amplitudes as a result of interindividual differences. Allowing the model classify such random variations in a random effects-term (1|ID) correctly models the within-subject character of the study.

Similarly, not only ID but also the prototype was included as a random intercept (1|Prototype). As prototypes elicit distinct ERP responses, reporting of respective effects would be trivial and, thus, fail to create new insight in terms of e.g. liking effects on ERP outcomes. In terms of random effects, random slopes were included throughout the analyses. For example, in the manipulation check analysis (see section *Results – Manipulation Check*), the design category variable was given a random slope as participants, arguably, increase differentially in liking moving from the *non-aesthetic* design category to the *aesthetic* design category.

EEG data of the study at hand were prepared using a custom-made Matlab 2022 preprocessing pipeline [32] – for further information consult the section *EEG Preprocessing and Data Analysis*. After preprocessing, the epoched ERP amplitudes were averaged over TWOI-1 and TWOI-2 and mean amplitudes were exported for further statistical analyses by means of mixed models in r-studio. Pre-processed ERP data also created the data base of the MVPA pattern analysis with MNE-python [36] and a contrast analysis was performed indicating predictions between the aesthetic and non-aesthetic category - answering the question whether the respective categories could be predicted above levels of chance. To explain decoding algorithm on the content-level, ERP-differences during both of the TWOIs were estimated by means of a mixed model per electrode and participant. As separate models were estimated per electrode and participant, the model Amplitude ∼ PrototypeCategory + (1 | Prototype) did not include the additional fixed and random effects of ROI, (1|electrode) or (1|ID). The resulting, model-derived estimated amplitude differences between stimulus categories per participant were, then, correlated with Haufe pattern values to provide a quantitative account of features underlying the decoding pipeline.

In order to assess inference statistical decoding accuracy significance, non-parametric permutation-based approaches were applied using functions from the MNE-Python toolbox [37] to control for multiple comparisons across time [48,49]. More specifically, at each time point, the decoding accuracy was compared against chance levels (50% accuracy) with one sampled *t*-tests. A cluster-based permutation approach was used to identify time-clusters and statistics were calculated as the sum of *t*-values within each cluster. Statistical significance was assessed by means of permutation testing – a process during which the sign of the observation was randomly flipped across participants to create a null-distribution of the maximum cluster-level statistics. Clusters were considered significant if their statistic exceeded the 95^th^ percentile of the null-distribution, effectively applying an alpha level of error probability of 5%. The respective procedures control for family-wise error rates while accounting for the fact that neighbouring timepoints are not independent in EEG data and, thereby, increasing sensitivity to lasting temporal effects [49,50]. By evaluating clusters of consecutive effects rather than isolated time points, the method increases sensitivity to temporally extended patterns of activity.

As cluster-based approaches evaluate the significance of clusters and not precise time points and locations [51], the method was complemented with an custom implementation of the maximum-statistics permutation approach [48,52] using SciPy [53] and NumPy [54]. A null-distribution was constructed by computing maximum the absolute t-value across all time points for each permutation. Observed *t*-values were then compared to this distribution to obtain family-wise error rate corrected *p*-values at each timepoint. Compared to the described cluster-based approach, this method is more conservative but it was applied to offer temporal specificity [50,52] about observed effects and, thus, allowed to identify peak decoding latencies.

In accordance with the hypothesis of an effect of individually liked and disliked NFB prototypes on ERP response and based on findings from logo design [7], TWOI-1 and TWOI-2 were defined as search spaces. Moreover, as earlier studies found significant effects of liking especially at central and parietal regions [7], the search was further limited to those locations. Analysis with respect to the parietal ROI were performed on ERP trajectories at the locations P3, Pz and P4 while results presented relating to central regions relate to the electrodes C3, Cz and C4.

To test our hypotheses regarding the effect of stimulus aesthetics on ERP response, different approaches to analyse data are possible. In previous studies [e.g. 7,55], participants rated all the stimuli regarding their visual appeal (liking measure) after stimulus exposure. Handy et al. [7], then, assessed, which 3 stimuli were liked the most and which three stimuli were liked the least. Comparisons were carried out between groups formed in accordance to the latter, categorical liking variable (liked, neutral, disliked).

This approach comes with some limitations. It can be expected that different stimuli produce specific ERP patterns as a result of their differences in terms of visual properties (e.g. the interaction between design characteristics such as colour, visual complexity, familiarity or concreteness). Combining individually liked and disliked stimuli for the ERP analysis hence implies that different stimuli are grouped together, which might lead to additional variances in ERP production (i.e. issue of comparing apples with pears).

The issue might be solved by another analysis approach using continuous measures of liking of all stimuli to predict their ERP responses. This procedure (termed ‘affective ex-post’ hereafter) focuses on individual subjective evaluations of liking of individual prototypes. By adding the random intercept term (1|Prototype) to the statistical model, predictor effects within prototypes can be statistically controlled for.

A further approach, circumventing the issue of uncontrolled variance in ERP response due to stimulus differences, is to analyse effects of liking within one specific prototype. The respective analysis will be termed ‘volitional ex-post’ hereafter referring to participants behavioral inclination of choosing a stimulus as their favourite prototype. Following this strategy, one of the 16 prototypes was chosen to analyse effects of liking on ERP responses. In order choose the ideal stimulus for the volitional ex-post approach, the following criteria were applied: (1) Maximum range of liking (range from the minimum - the maximum of the scale), (2) the group sizes of a given prototype being chosen as Favourite / Neither / Least-Favourite should be >5 to ensure reasonable group sizes, and finally, (3) the standard deviation of liking should be as large as possible.

A respective data analysis showed that criterion 1was met by prototypes 1, 2, 3, 4, 5, 7, 8, 9, 10, 11, 14 and 16. Of those, only prototypes 2, 3 and 4 met criterion 2 (see Fig 6, panels c and d), requiring a rather equal distribution regarding the rating for the most or least favourite prototype. In terms of variance, prototypes 2, 3 and 4 evoked standard deviations of 1.68, 1.97 and 1.55, respectively. Prototype 3, obtaining the highest variance regarding ratings of liking, was selected (see Fig 4, panel a) for the analysis following the ‘volitional ex-post’ approach.

**Fig 6.**
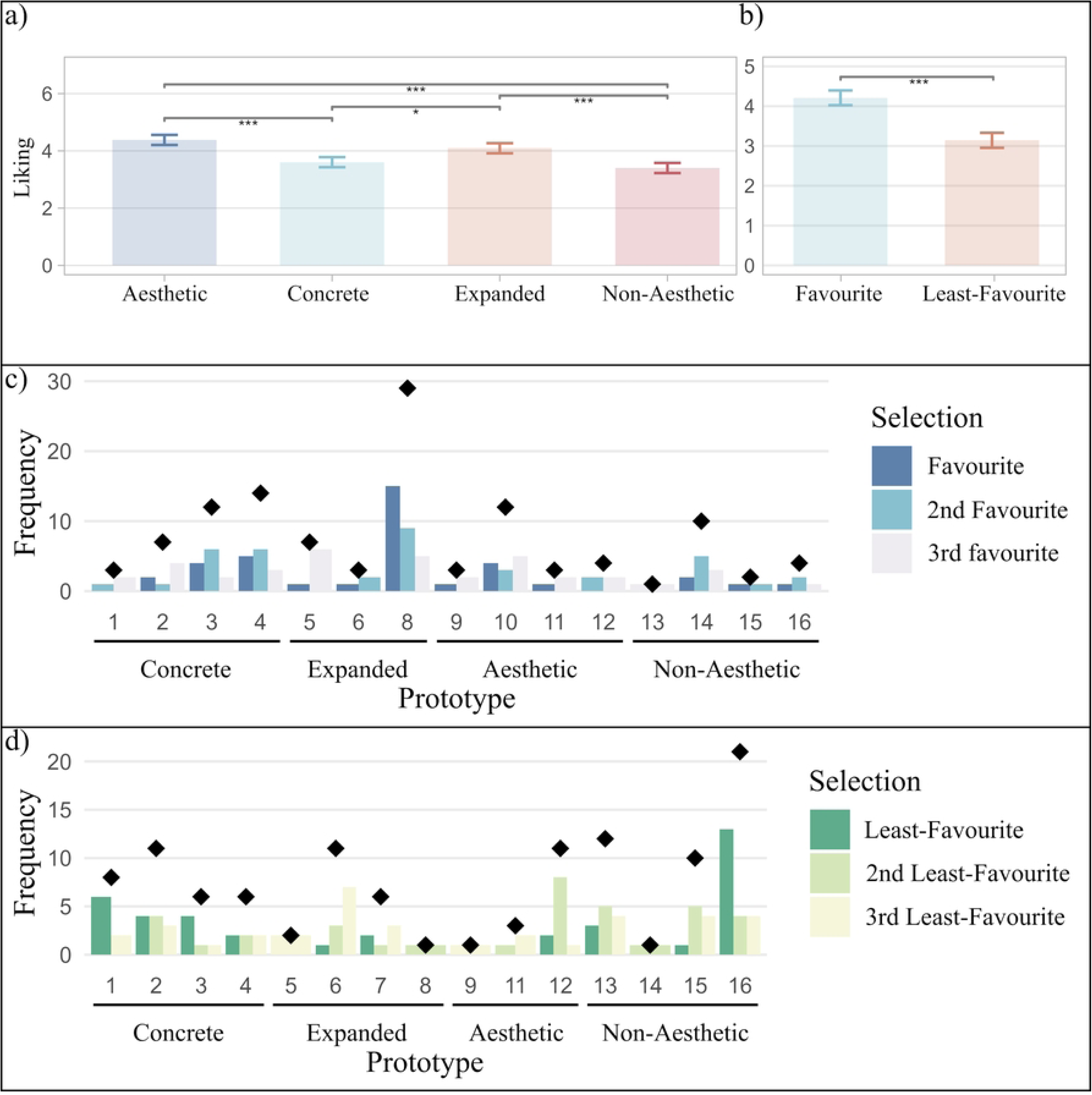
Liking and Preference of NFB Stimuli. Panel a and b show liking in response to different stimulus categories – each of the categories comprised four neurofeedback training design stimuli. Panel b depicts liking in response to preference. The favourite category includes the three prototypes individually selected as favourite, second favourite or third favourite prototype. Similarly, the least-favourite category includes prototypes individually chosen as least-favourite, second least-favourite or third least-favourite prototype. Panel a and b error bars constitute standard error of the mean (*SEM*). Panel a and b error probabilities were corrected for multiple comparisons by means of “fdr”. Level of significance indicated by means of asterisks: *p* < .05’, *p* < .01*, *p* < .001**, *p* < .0001***. Panel c depicts frequencies of preference per prototype. Individual prototypes selected as favourites are printed blue and least-favourite selections are shown in green (panel c). Panel c and d bars depict the frequency a prototype was mentioned as (least) favourite, second (least) favourite or third (least) favourite. Sum values of preference are indicated by diamond shapes.

Finally, a third approach was developed to test our hypotheses (termed ‘*categorical ex-ante*’) adopting a controlled experimental procedure. Following this approach, design aesthetics of stimuli were manipulated experimentally referring to specific design principles and guidelines (c.f. section on *Stimuli* above). Controlling for visual properties of the aesthetically pleasing and displeasing stimuli by design, the two sets of prototypes are compared regarding their EPR responses including the critical error term (1|Prototype).

## Results

### Manipulation check

In order to check whether the design manipulation was successful a random intercept model defined as Liking ∼ Category + (1|ID). The model improved fit in comparison to model 0, *X*^2^(3, *N* = 38) = 43.22, *p* < .001. The categories differed with respect to liking, *F*(3, 567) = 14.89, *p* < .001, η ^2^ = .073, see Fig 6, panel a.

Contrasts revealed that the category non-aesthetic was liked the least (*EMM* = 3.30, 95% *CI* [3.03, 3.58]) while the Aesthetic category was liked the most (*EMM* = 4.27, 95% *CI* [3.96, 4.58]). Category differences are depicted in Table 1.

**Table 1.** Contrasts of Estimated Means of Liking.

| Contrast | $\Delta$ | *SEM* | *df* | *t* | *p* | $\eta_p^2$ |
| --- | --- | --- | --- | --- | --- | --- |
| Non-Aesthetic vs. Concrete | 0.20 | 0.16 | 567 | 1.24 | 1.000 | .003 |
| Non-Aesthetic vs. Expanded | 0.69 | 0.16 | 567 | 4.21 | <.001 | .030 |
| Non-Aesthetic vs. Aesthetic | 0.98 | 0.16 | 567 | 5.98 | <.001 | .059 |
| Concrete vs. Expanded | 0.49 | 0.16 | 567 | 2.97 | .019 | .015 |
| Concrete vs. Aesthetic | 0.78 | 0.16 | 567 | 4.73 | <.001 | .038 |
| Expanded vs. Aesthetic | 0.29 | 0.16 | 567 | 1.77 | .469 | .006 |

*Abbreviations:* *df*- degrees of freedom; *p* - probability of error, *SEM* - standard error of the mean. *T*-tests were corrected for multiple comparisons by means of Bonferroni (factor 6).

Similarly to liking and complexity, the four stimulus categories differed in terms of preference.

Frequencies of (least) favourite prototypes, second (least) favourite prototypes and third (least) favourite prototype can be consulted in Fig 6, panels c and d.

As aesthetic evaluation lies in the eye of the beholder (e.g. Silvia, 2005), liking ratings were analysed as a function of preference (see Fig 6, panel b). As expected, the three favourite stimuli obtained higher liking ratings (*EMM* = 4.21, 95% *CI* [3.84, 4.58]) than the three least-favourite stimuli (*EMM* = 3.14, 95% *CI* [2.77, 3.52]) and the difference was significant, *b* = 1.07, *SE* = 0.21, *t*(185.32) = 5.11, *p* < .001, η² = .12.

### ERP in response to liking & complexity ratings (affective ex-post approach)

The random intercept, random slope models for amplitudes at TWOI-1 and TWOI-2 were defined in the following way accounting for random effects of prototype (1|Prototype) and participants (1|ID) to model the within subject and within prototype design:

Amplitude ∼ Liking * Complexity + ROI + Liking : ROI + Complexity : ROI + (1 + Complexity * Liking || ID) + (1 | Prototype) + (1 | Electrode)

For both TWOI-1 (*X*^2^ (6, *N* = 38) = 23.30, *p* < .001) and TWOI-2 (*X*^2^ (6, *N* = 38) = 49.20, *p* < .001), the model improved fit in comparison to model 0. Significant effects were found with respect to the ERP response predicted by beholder-based liking and complexity as well as their interaction (see Table 2, Table 3, and Fig. 7). At both TWOIs, results were in accordance with our hypotheses - liking NFB affected ERP responses independently from individual prototypes and perceiving a stimulus as complex showed similar effects. Note, none of the effects interacted with ROI indicating both regions contributed to the process.

**Fig 7.**
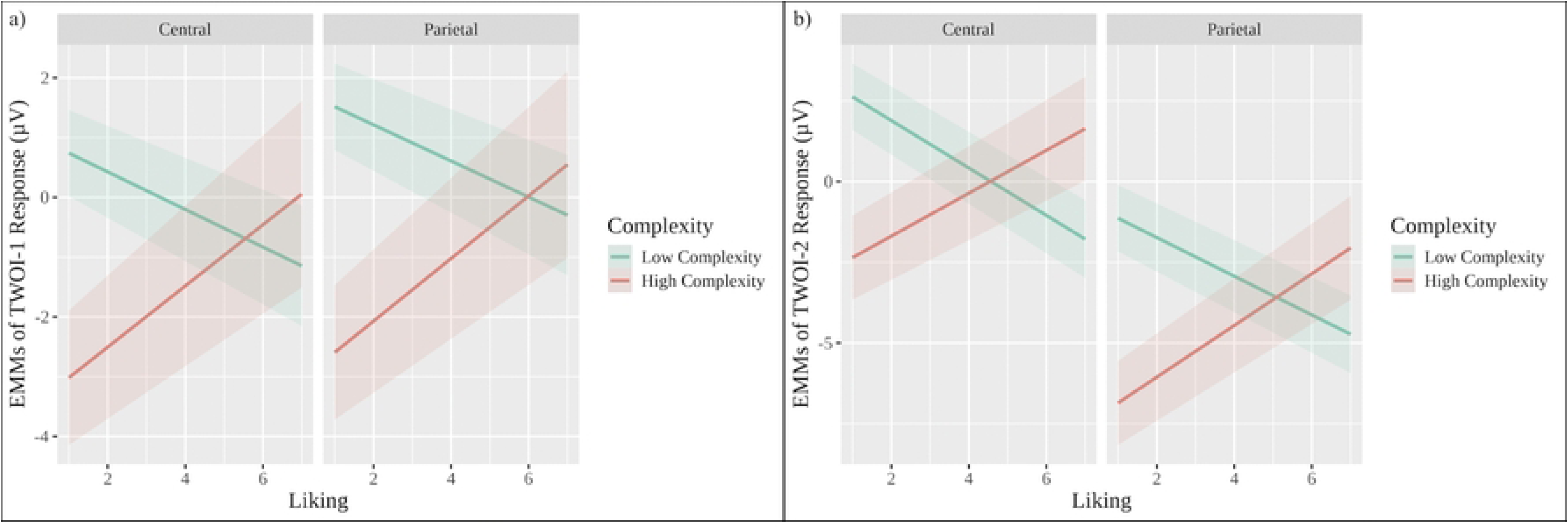
ERP Patterns in Response to Liking and Complexity. Panels a and b depict TWOI-1 and TWOI-2 in response to liking, complexity. TWOI-1 was defined between 100-200 ms while TWOI-2 was defined between 200-300. The central ROI included C3, Cz and C4, the parietal ROI included P3, Pz and P4. Note, stimuli presentation took place at time point 0, electrode positions were collected according to the international 10-20 system. The baseline time (from -100 to 0 ms seconds of stimulus presentation) was set to 0. Ribbons indicate *SEMs*. Abbreviations in the graph are event-related potential (ERP), time window of interest (TWOI) and region of interest (ROI).

**Table 2.**
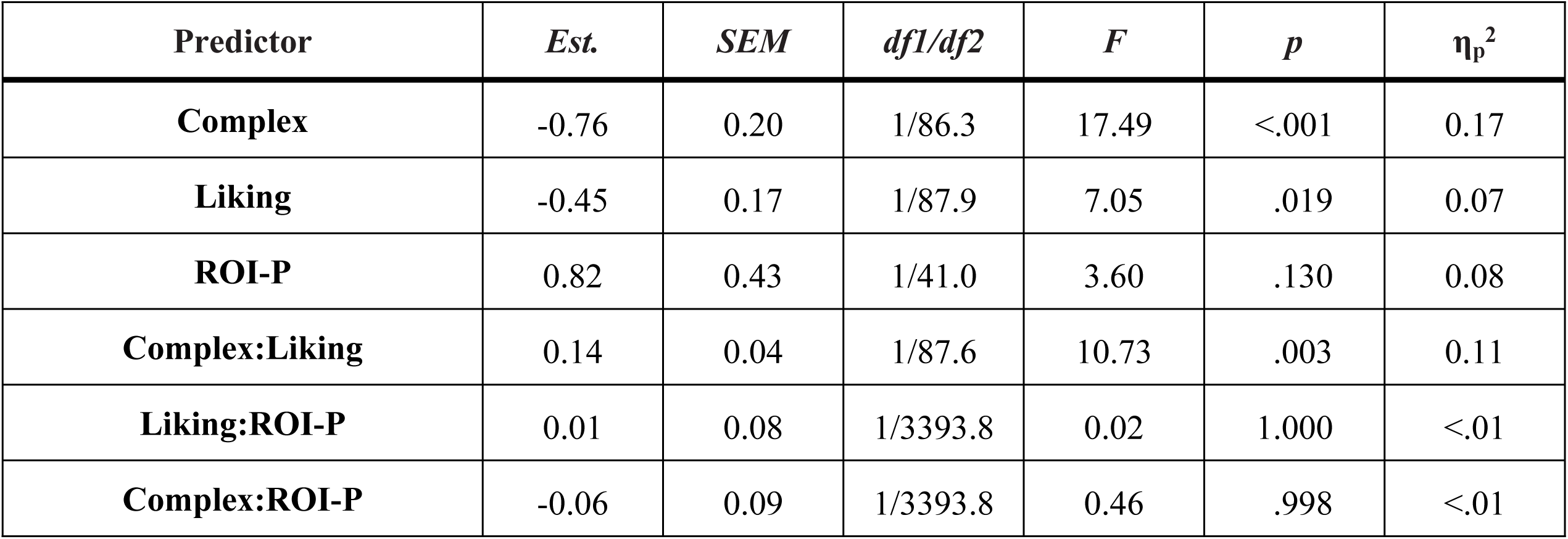
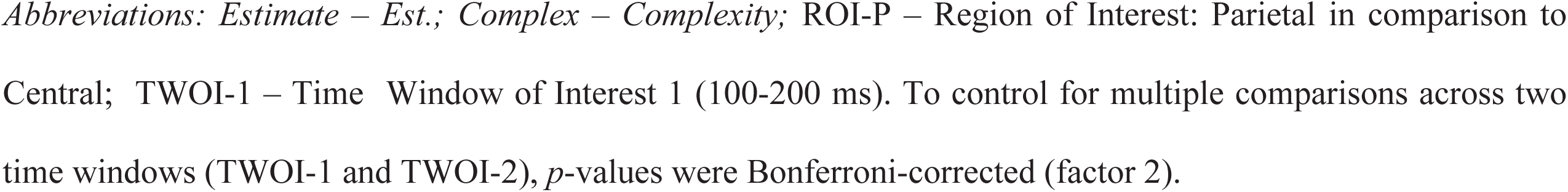
Mixed-Model Regression Analyses: Effects of Predictors Liking, Complexity and ROI on TWOI-1 Response.

| Predictor | *Est.* | *SEM* | *df1/df2* | *F* | *p* | $\eta_p^2$ |
| --- | --- | --- | --- | --- | --- | --- |
| **Complex** | -0.76 | 0.20 | 1/86.3 | 17.49 | <.001 | 0.17 |
| **Liking** | -0.45 | 0.17 | 1/87.9 | 7.05 | .019 | 0.07 |
| **ROI-P** | 0.82 | 0.43 | 1/41.0 | 3.60 | .130 | 0.08 |
| **Complex:Liking** | 0.14 | 0.04 | 1/87.6 | 10.73 | .003 | 0.11 |
| **Liking:ROI-P** | 0.01 | 0.08 | 1/3393.8 | 0.02 | 1.000 | <.01 |
| **Complex:ROI-P** | -0.06 | 0.09 | 1/3393.8 | 0.46 | .998 | <.01 |

Abbreviations: *Estimate* – *Est.*; *Complex* – *Complexity*; ROI-P – Region of Interest: Parietal in comparison to Central; TWOI-1 – Time Window of Interest 1 (100-200 ms). To control for multiple comparisons across two time windows (TWOI-1 and TWOI-2), *p*-values were Bonferroni-corrected (factor 2).

**Table 3.** Mixed-Model Regression Analyses. Effects of Predictors Liking, Complexity and ROI on TWOI-2 response (200-300 ms)

| Predictor | *Est.* | *SEM* | *df1/df2* | *F* | *p* | $\eta_p^2$ |
| --- | --- | --- | --- | --- | --- | --- |
| **Complex** | -1.06 | 0.20 | 1/127.75 | 33.15 | <.001 | .20 |
| **Liking** | -0.97 | 0.18 | 1/124.21 | 26.99 | <.001 | .17 |
| **ROI-P** | -3.77 | 0.90 | 1/6.37 | 17.39 | .010 | .02 |
| **Complex:Liking** | 0.23 | 0.04 | 1/128.62 | 27.72 | <.001 | .17 |
| **Liking:ROI-P** | 0.14 | 0.09 | 1/3441.74 | 2.13 | .289 | <.01 |
| **Complex:ROI-P** | -0.12 | 0.10 | 1/3441.74 | 1.53 | .432 | <.01 |

Abbreviations: Estimate – Est.; Complex – Complexity; ROI-P – Region of Interest: Parietal in comparison to Central; TWOI-2 – Time Window of Interest 2 (200-300 ms). To control for multiple comparisons across two time windows (TWOI-1 and TWOI-2), p-values were Bonferroni-corrected (factor 2).

The obtained interaction effect between liking and complexity was further explored with a trend analyses (multiple comparison correction across TWOIs, factor 2). Results showed a non-significant negative trend effect of liking on ERP amplitudes, when complexity was low (*b_LowComplexity_* = -0.31, 95% *CI* [-0.59, 0.02], *t*(73.20) = 2.15, *p* = 0.069) but a significant positive effect was found when complexity was high (*b_HighComplexity_* = 0.52, 95% *CI* [-0.09, 0.95], *t*(51.70) = 2.40, *p* = 0.040). Additionally, the slope of liking on TWOI-1 amplitude was significantly more positive when complexity was high Δ*b_High-LowComplexity_*= 0.83, 95% *CI* [0.32, 1.33], *t*(121.00) = 3.22, *p* = 0.003.

Effect sizes were descriptively smaller for TWOI-1 in comparison to TWOI-2 (see Table 3). For example, the effect of liking at TWOI-1 (η_p_ ^2^ = .07) can be considered a medium-sized effect [56] while liking shows a large effect at TWOI-2 (η_p_ ^2^ = .17).

Exploring the interaction liking:complexity, follow-up trend analyses (applying Bonferroni correction, factor 2) indicated a significant negative effect of liking in TWOI-2 amplitude when complexity was low (*b_LowComplexity_* = -0.67, 95% *CI* [-0.95, 4.63], *t*(85.30) = -4.63, *p* = <.001). Moreover, a significant positive effect was found when complexity was high (*b_HighComplexity_* = 0.73, 95% *CI* [0.31, 1.16], *t*(57.00) = 3.45, *p* = 0.002). Finally, the slopes differed significantly and the effect of liking on TWOI-2 amplitude was more positive when complexity was high Δ*b_High-LowComplexity_* = 1.40, 95% *CI* [0.86, 1.94], *t*(157.00) = 5.16, *p* < .001.

### ERP in response to preference (volitional ex-post approach)

As the data set for within prototype analysis was smaller and comprised 1/16th of the data, the applied model was simplified. Consecutively, possible effects of Preference (Least-Favourites = -1; Neutral = 0; Favourites = 1) at TWOI-1 and TWOI-2 were evaluated (see Fig 8 and Table 4) with the following random intercept model:

**Fig 8.**
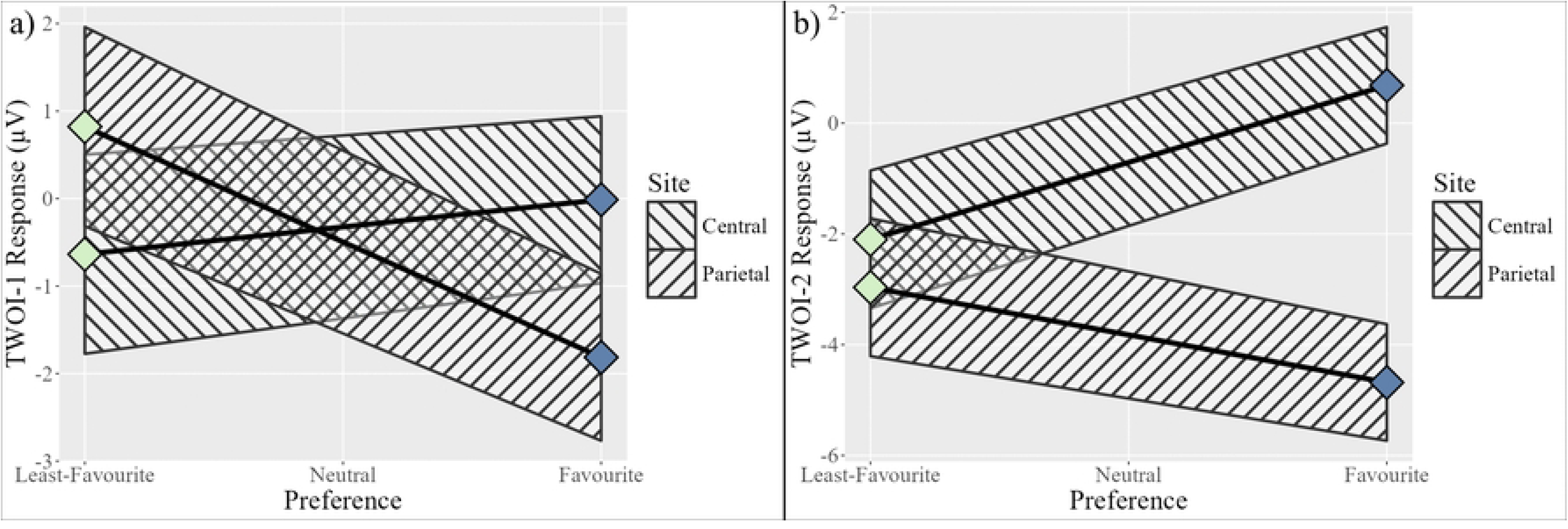
ERPs in response to NFB Prototype Preference at TWOI-1 (panel a) and TWOI-2 (panel b). NFB-Prototype categories “Favourite”, “Least favourite” and “Neutral” represent the number of times prototype 3 was selected as a favourite, a least-favourite or was not mentioned as either and, thus, classified as “neutral”. Note, the central ROI included C3, Cz and C4, the parietal ROI included P3, Pz and P4, stimuli presentation took place at time point 0, electrode positions were collected according to the international 10-20 system. The baseline time (from -100 to 0 ms seconds of stimulus presentation) was set to 0. Ribbons indicate *SEMs*. Abbreviations in the graph are event-related potential (ERP), time window of interest (TWOI) and region of interest (ROI).

**Table 4.** Mixed-Model Analyses: Effects of Predictors Preference and ROI on TWOI-1 (100-200 ms) and TWOI-2 (200-300 ms) Response.

| **TWOI-1** | *Est.* | *SEM* | *df1/df2* | *F* | *p* | $\eta_p^2$ |
| --- | --- | --- | --- | --- | --- | --- |
| **Pref** | 0.31 | 0.86 | 1/36.00 | 0.36 | 1.000 | 0.01 |
| **ROI-P** | -0.17 | 0.37 | 1/4.13 | 0.22 | 1.000 | 0.05 |
| **Pref:ROI-P** | -1.63 | 0.35 | 1/184.00 | 21.45 | <.001 | 0.10 |
| **TWOI-2** | *Est.* | *SEM* | *df1/df2* | *F* | *p* | $\eta_p^2$ |
| **Pref** | 1.39 | 0.91 | 1/36.00 | 0.09 | 1.000 | <.01 |
| **ROI-P** | -3.12 | 0.55 | 1/4.08 | 31.45 | .009 | 0.89 |
| **Pref:ROI-P** | -2.25 | 0.42 | 1/184.00 | 29.30 | <.001 | 0.14 |

*Abbreviations:* ROI-P – Region of Interest - parietal in comparison to central. To control for multiple comparisons across two time windows (TWOI-1 and TWOI-2), *p*-values were Bonferroni-corrected (factor 2).

Amplitude ∼ Preference * ROI + (1|ID) + (1 | Electrode)

The models for TWOI-1, *X*^2^ (3, *N* = 38) = 22.05, *p* < .001; and TWOI-2, *X*^2^ (3, *N* = 38) = 40.10, *p* < .001; improved fit in comparison to model 0. Results indicated no main effect of preference during TWOI-1 and TWOI-2. However, analysis revealed a main effect of ROI during TWOI-2 indicating different amplitudes at the parietal ROI in comparison to the central ROI (see Table 4). Moreover, an interaction effect between ROI and preference was observed both, during TWOI-1 and TWOI-2 - speaking in favour of altered phasic EEG activity in response to preference. Exploration of interaction effects (see Fig 8) revealed a disordinal interaction for TWOI-1 and an ordinal interaction during TWOI-2.

Follow-up trend analyses of estimated marginal means were calculated applying multiple comparison control with Bonferroni adjustment across TWOIs (factor 2). Results showed that the relationship between preference and TWOI-2 amplitudes was not significant at either ROI (*b_Central_*= 0.31, 95% *CI* [-1.42, 2.05], *t*(39.2) = 0.36, *p* = 1.000; *b_parietal_* = -1.32, 95% *CI* [-3.05, 0.42], *t*(39.2) = - 1.54, *p* = 0.264. However, slopes differed significantly between the central and the parietal ROIs, Δ*b_central-parietal_*= 1.63, 95% *CI* [0.94, 2.32], *t*(184) = 4.63, *p* < .001. The result indicates that preference had a more positive effect on central amplitudes vs. on parietal amplitudes.

Similarly, during TWOI-2, individual slopes describing the connection between preference and TWOI-2 amplitudes were not significant (*b_Central_* = 1.40, 95% *CI* [-0.46, 3.24], *t*(40) = 1.52, *p* = 0.272; *b_parietal_* = -0.86, 95% *CI* [-2.71, 0.99], *t*(40) = -0.94, *p* = 0.707). Again, a significant difference was found between slopes at the central and the parietal ROI; Δ*b_central-parietal_* = 2.25, 95% *CI* [1.43, 3.07], *t*(184) = 5.41, *p* < .001. Thus, the relationship between preference and amplitudes was significantly more positive at the central ROI in comparison to the parietal ROI.

### ERPs in response to manipulated prototype categories (categorical ex-ante approach)

Hypothesising that neural responses at TWOI-1 and TWOI-2 are similar in response to manipulated prototype categories (aesthetic vs. non-aesthetic) to beholder-based liked vs. disliked stimuli, ERP responses were analysed. Although the visual inspection of EPR response graphs seemed to indicate different trajectories in response to the manipulated prototype categories (see Fig 9 and supplementary material b), those differences did not persist, when the prototype variable was considered in the statistical random intercept, random slope models for TWOI-1 and TWOI-2 (see Table 5):

**Fig 9.**
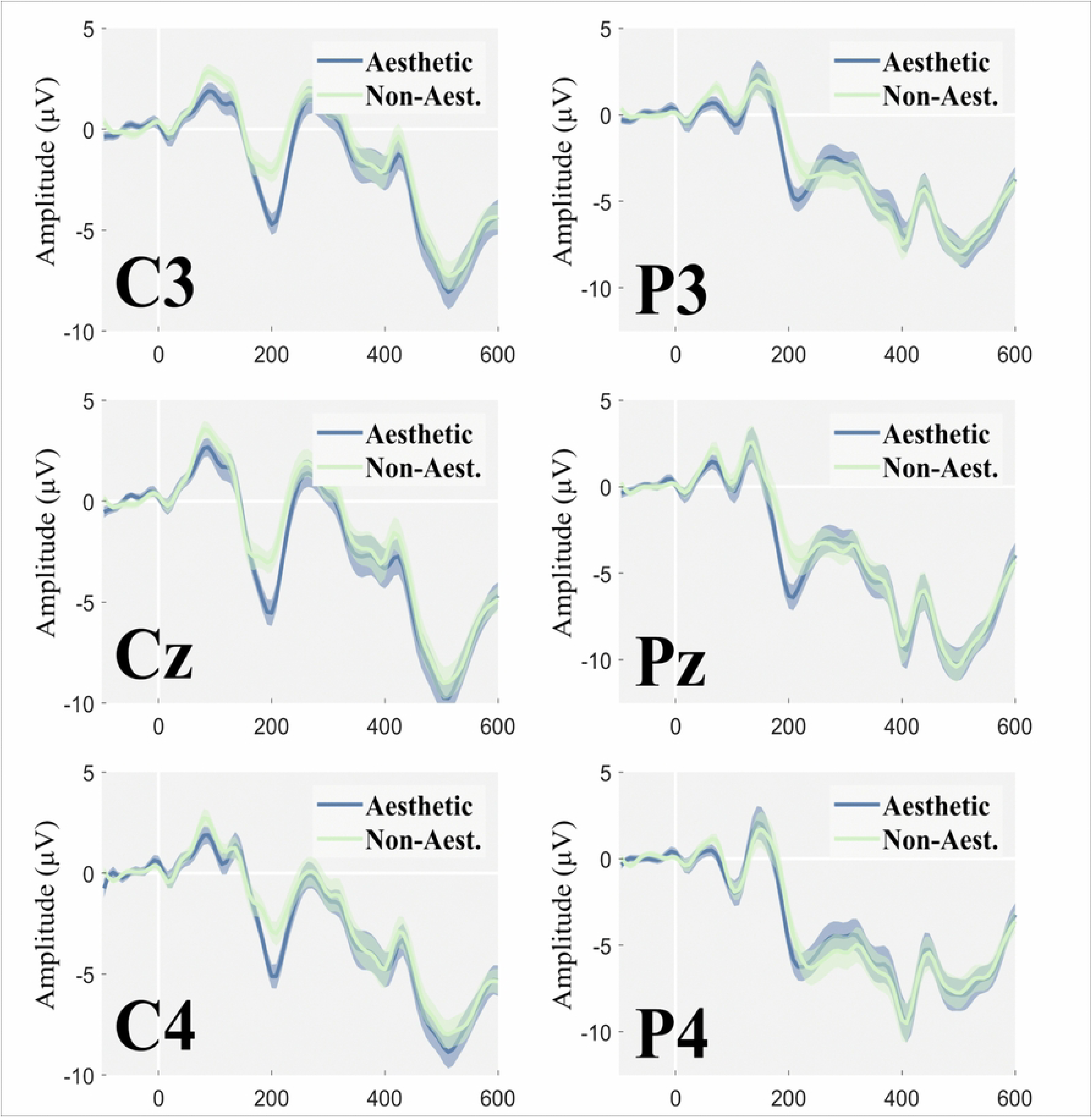
ERPs in Response to Manipulated Prototype Categories “Aesthetic” vs. “Non-Aesthetic”. Note, stimuli presentation took place at time point 0, EEG electrodes were placed according to internation 10-20 system. The baseline time (from -100 ms to 0 ms seconds of stimulus presentation) was set to 0. Shaded areas around the curve represent *SEMs*. Abbreviations in the graph are event-related potential (ERP).

**Table 5.** Mixed-Model Analyses Regression: Effects of Manipulated Prototype Category (Aesthetic vs. Non-Aesthetic) and ROI on TWOI-1 (100-200 ms) and TWOI-2 (200-300 ms)

| TWOI-1* | *Est.* | *SEM* | *df1/df2* | *F* | *p* | $\eta_p^2$ |
| --- | --- | --- | --- | --- | --- | --- |
| ProtoCat | -0.74 | 1.06 | 1/6.39 | 0.37 | 1.000 | 0.05 |
| ROI-P | 0.51 | 0.37 | 1/4.00 | 3.61 | .260 | 0.47 |
| **ProtoCat:ROI** | 0.19 | 0.38 | 1/1736.00 | 0.26 | 1.000 | <.01 |
| **TWOI-2** | *Est.* | *SEM* | *df1/df2* | *F* | *p* | $\eta_p^2$ |
| **ProtoCat** | -1.17 | 0.85 | 1/6.39 | 0.85 | .776 | 0.11 |
| **ROI-P** | -3.85 | 0.85 | 1/4.00 | 17.65 | .028 | 0.82 |
| **ProtoCat:ROI** | 0.81 | 0.40 | 1/1736.00 | 3.96 | .094 | <0.01 |

*Abbreviations:* ProtoCat - Prototype Category; ROI-P – Region of Interest – Parietal. To control for multiple comparisons across two time windows (TWOI-1 and TWOI-2), *p*-values were Bonferroni-corrected (factor 2). Note, for TWOI-1\*, adding the predictors did not significantly improve model fit relative to the model 0; results are included for completeness solely.

Amplitude ∼ PrototypeCategory * ROI + (1 + PrototypeCategory | ID) + (1|Electrode) + (1|Prototype)

For TWOI-1, the model did not improve fit in comparison to model 0, *X*^2^ (3, *N* = 38) = 3.97, *p* = 0.265 and therefore TWOI-1 analyses are presented solely for completeness. Comparing the TWOI-2 model to model 0 improved fit significantly, *X*^2^ (3, *N* = 38) = 14.08, *p* = 0.003, see Table 5 and Fig 9. For TWOI-2, the respective random intercept, random slope model did not indicate any main effects of prototype category or interaction effects with ROI after results were corrected for multiple comparisons by means of Bonferroni.

### ERP-Decoding

In order to increase convergent validity, ERPs were analysed by means of MVPA decoding [35]. More specifically, the decoding pipeline predicted manipulated prototype categories (aesthetic vs. non-aesthetic) by means of ERPs at the central and parietal ROIs (see Fig 10, panel a).

**Fig 10.**
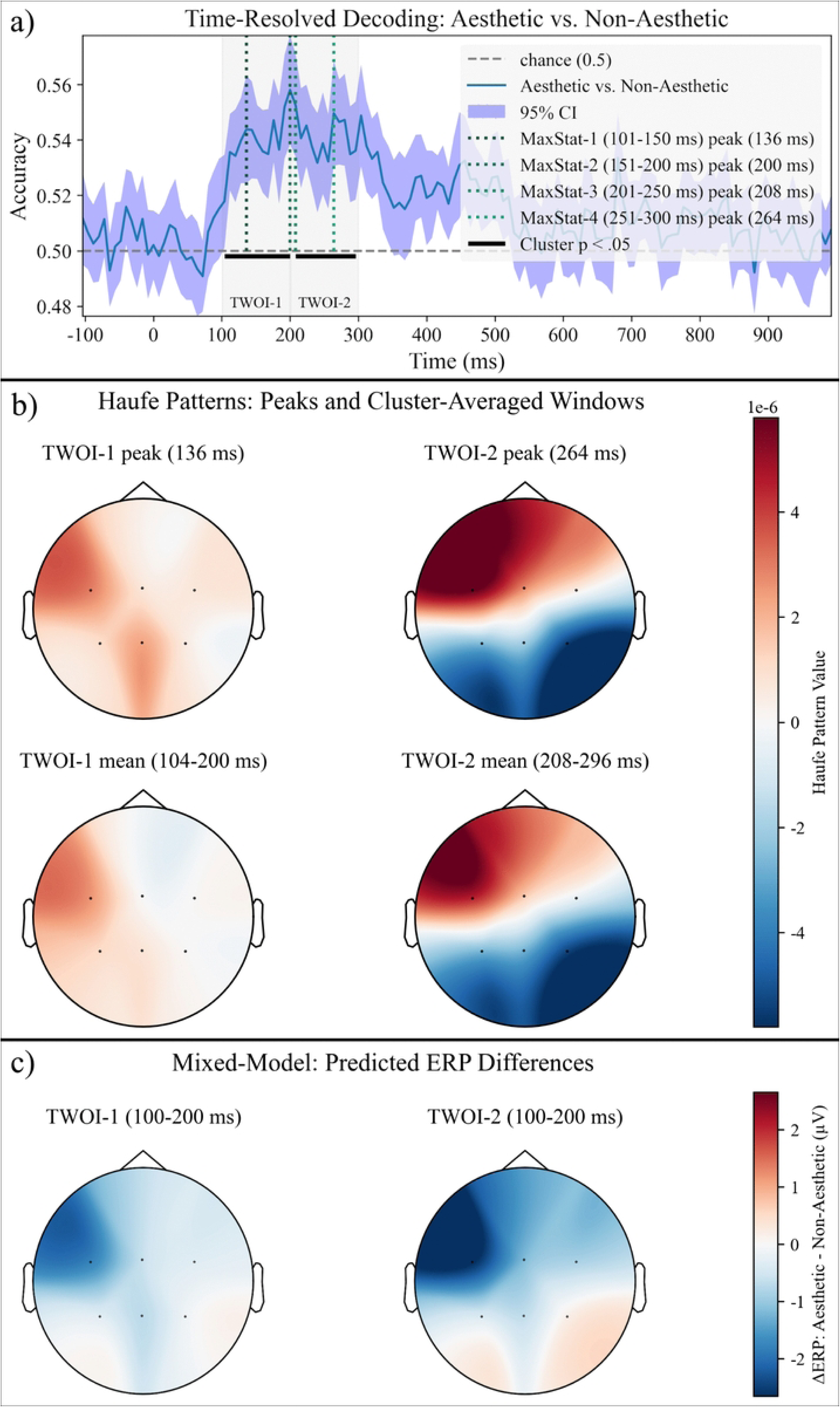
Decoding Accuracy in Response to Manipulated Prototype Categories “Aesthetic” vs “Non-Aesthetic”. Panel a shows decoding accuracy in percent as well as significant peaks and time windows during TWOI-1 and TWOI-2. Panel b topographical Haufe patterns derived by the decoding pipeline for the Categories “Aesthetic” and “Non-Aesthetic”. Panel c depicts mixed model category differences defined as aesthetic - non-aesthetic. Stimulus presentation took place at time point 0, electrode positions were collected according to the international 10-20 system and shaded areas around the curve depicting 95% *CI*. Abbreviations in the graph are event-related potential (ERP).

The confirmatory decoding inference statistics within TWOI-1 by cluster-based permutation indicated a time window of 104-200 ms significantly differing from chance (*M_AccuracyTwoi1_* = 53.99% (95% *CI* [53.01, 54.97]*, p* <.001). Maximum-statistics permutation approach indicated that the peak accuracies within TWOI-1 occurred after 136 ms and 200 ms differing significantly from chance; *Max_AccuracyTwoi1-136_* = 54.38%, 95% CI [52.44, 56.32], *t*(37) = 4.43, *p* = 0.001, *d* = 0.72; *Max_AccuracyTwoi1- 200_* = 55.81%, 95% CI [53.85, 57.77], *t*(37) = 5.80, *p* <.001, *d* = 0.94. Similarly, for TWOI-2, according to the cluster-based permutation, within the time window of 200-296 ms, prediction accuracy differed from chance (*M_AccuracyTwoi2_* = 54.11% (95% *CI* [53.08, 55.14]*, p* <.001). During TWOI-2, the maximum-statistics permutation approach pointed at peak prediction accuracies after 208 ms and 264 ms; *Max_AccuracyTwoi2-208_* = 55.26%, 95% *CI* [53.40, 57.13], *t*(37) = 5.53, *p* <.001, *d* = 0.90; *Max_AccuracyTwoi2-264_* = 54.96%, 95% *CI* [53.06, 56.86], *t*(37) = 5.12, *p* <.001, *d* = 0.83.

The overlap of predicted ERP differences (see Fig 10c) and Haufe patterns [see Fig 10b, 57] was indicated by means of Bonferroni corrected Pearson correlations during TWOI-1 and TWOI-2, see Table 6. These correlational results (see Fig 11) suggest that the decoding patterns are closely linked to ERP amplitude variations at the a priori defined ROIs and TWOIs – suggesting that the classifier relies on interpretable, physiologically meaningful signal differences.

**Fig 11.**
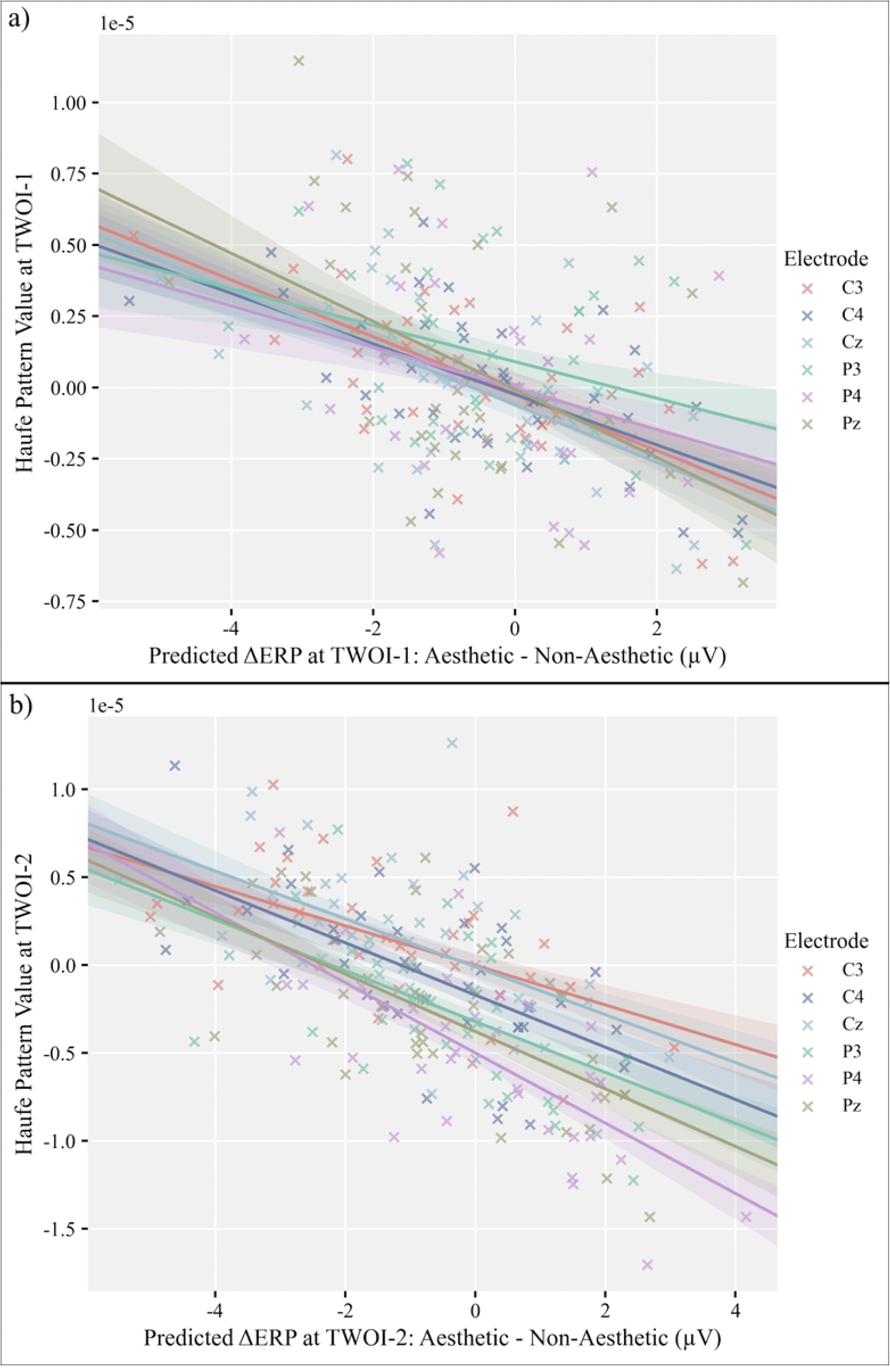
Correlations of Predicted ERP Differences and Haufe Patterns During TWOIs. Panel a shows correlations during TWOI-1, panel b depicts correlations during TWOI-2. Electrode positions were collected according to the international 10-20 system and shaded areas around the curve depicting *SEMs*. Abbreviations in the graph are event-related potential (ERP) and time window of interest (TWOI).

**Table 6.** Correlations between predicted ERP differences at ROI electrodes during TWOI-1 and TWOI-2, respectively. Error probabilities controlled for multiple comparisons by means of Bonferroni per TWOI.

| TWOI-1 | *r* | *df* | *t* | *p* | *R*<sup>2</sup> |
| --- | --- | --- | --- | --- | --- |
| C3 | -0.62 | 35 | -4.72 | <.001 | 0.39 |
| C4 | -0.61 | 36 | -4.59 | <.001 | 0.37 |
| Cz | -0.54 | 36 | -3.86 | .003 | 0.29 |
| P3 | -0.28 | 36 | -1.77 | .508 | 0.08 |
| P4 | -0.31 | 36 | -1.97 | .338 | 0.10 |
| Pz | -0.47 | 36 | -3.18 | .018 | 0.22 |
| TWOI-2 | *r* | *df* | *t* | *p* | *R*<sup>2</sup> |
| C3 | -0.54 | 35 | -3.79 | .003 | 0.29 |
| C4 | -0.62 | 36 | -4.78 | <.001 | 0.39 |
| Cz | -0.57 | 36 | -4.13 | .001 | 0.32 |
| P3 | -0.56 | 36 | -4.08 | .001 | 0.32 |
| P4 | -0.68 | 36 | -5.53 | <.001 | 0.46 |
| Pz | -0.64 | 36 | -5.05 | <.001 | 0.42 |

## Discussion

The aim of the study at hand was two-fold. On the one hand, it was evaluated whether the application of design principles would show a respective neural correlate assessed by means of ERP analysis, indicating systematic variations in cortical processing associated with the experimental manipulation. On the other hand, it was explored whether results from the context of visual design could be replicated with NFB stimuli. Specifically, the study addressed the question whether liking NFB stimuli and perceiving them as complex affected ERP trajectories, independently of presumable effects of visual differences of NFB prototypes. Indeed, respective effects were found at central and parietal ROIs.

In contrast to previous ERP studies on visual design aesthetics, the current study employed three complementary analytical approaches to address methodological limitations, particularly the fixed-effects fallacy, to strengthen the convergent reliability of the findings. Two approaches analysing within prototype effects of liking and preference on TWOI response were deployed, see results of the *affective ex-post* approach and the *volitional ex-post* approach. Both methods yielded similar results - the volitional ex-post approach showed increased N1 and N2 amplitudes as a result of preference at the parietal ROI during both TWOIs (see Fig 8). This result was replicated by the affective ex-post approach showing a negative main effect of liking on ERP amplitudes during both TWOIs at both ROIs (see Table 2 and Table 3).

Exploring the negative effect of liking on TWOI-2 could be interpreted in terms of processing fluency theory [58] as ease of processing has a favourable effect on the perceiver’s aesthetic reaction. More specifically, a less pronounced P2 component (see Fig 7, panel b) can be interpreted as an indicator of fluent processing as ERP amplitudes can be conceptualised in terms of the amount of neural resources participating during information processing [59].

Interpreting obtained effects (see Fig 7), it can be conjectured that the following pattern is observed: Smaller component deflection result from liked stimuli, potentially indicating improved processing fluency. Importantly liking interacted with perceived complexity (see Table 2 and Table 3) – an effect that is in line with findings indicating that some perceivers prefer complex stimuli while other prefer simple stimuli [60]. Complex, disliked stimuli produced a more pronounced N1 at central locations and a more pronounced N2 at parietal locations in comparison to disliked simple stimuli (see Fig 7). Additionally, N1/P1 component deflections were smaller for liked stimuli (see Fig 7, panel a) speaking in favour of ease of processing the stimuli. Interestingly, N1/P1 deflections increased for disliked stimuli, potentially indicating processing difficulty further strengthening the processing fluency interpretation.

The obtained interaction effects between liking and complexity can also be interpreted following the Integrative Theory of Aesthetic Value by Brielmann and Dayan [61] merging processing Fluency Theory and Learning Theories [see e.g., 62]. The theoretical frame-work of learning theories somewhat conflicts with processing fluency. Processing fluency posits that easily processed stimuli (i.e. simple stimuli) lead to more aesthetically pleasing experiences [8]. Learning theories, however, emphasize the formation of an internal cognitive model as a central mechanism for optimising the processing of future experiences. In other words, *adequate* stimuli should be more aesthetically pleasing – stimuli which are not necessarily simple.

Brielmann and Dayan’s approach integrates those two lines of argument by defining aesthetic value as a result of “any immediate reward – the first element of the long-run future – with the expectation of the integrated value of the second and subsequent elements” [61]. Or, simplified, two mechanisms are at work. The first process includes a positive effect of simple stimuli, increasing processing fluency adding to the aesthetic value. The second process of aesthetic perception involves the integral process of cognitive systems continuously predicting future events and adapting the prediction model. Stimuli deviating from the future prediction are aesthetically pleasing, when general uncertainty is low [63]. Applying Brielman and Dayan’s theory top the current results, aesthetic value might be diminished if simple stimuli strongly deviate from the internal model and, thus, integrating them potentially impinges on future predictions. The sum evaluation of both, the expectation of the usefulness of integrating this stimulus into the internal model and the immediate reward i.e. processing fluency results in the so-called *affective prediction error*.

Indeed, the affective prediction error seems to explain the observed P1/N1 and P2/N2 result pattern. Exploring simple stimuli effects according to Processing Fluency Theory, simplicity should have a mere positive effect on liking. This is true for some conditions resulting in exhibiting less pronounced P1/N1 component activation at both ROIs. However, some simple stimuli were disliked and led to larger P1/N1 as well as P2/N2 activation. This effect pattern (see Fig 7) does not converge with Processing Fluency Theory. However, this finding may indicate that simple stimuli did not substantially contribute to updating the internal predictive model, particularly in the context of the distraction task (scrambles vs. NFB prototype). Or, comparably higher complexity was more representative for the task at hand, and thus, was favoured by the internal model and positively reinforced by ease of processing and, less pronounced component activation.

For the complete interpretation TWOI-2 component activation patterns it seems crucial to discuss the increased P2/N2 activation for peak liking conditions. This result aligns with previous findings [64,65] indicating increased P2 amplitudes as a function of evaluative intensity. Accordingly, for perceivers with a preference for complex stimuli [60], complex stimuli might have elicited a stronger P2 reflecting the stronger positive appraisal. Additionally, larger N2s have also been linked to the arousal levels of both positive and negative pictures [66] indicating that increased N2 of liked simple stimuli might reflect the stronger positive response of participants preferring simple stimuli.

Finally, the disordinal interaction effect between liking and complexity affecting could be connected to dynamics of repeated encounters in the Integrative Theory of Aesthetic Value [61]. When a stimulus is encountered repeatedly liking increases due to mere exposure effects and, after a certain amount of encounters re-decreases as a result of habituation/boredom forming an inverted U-shape curve [25]. The effects of repetitive stimulus exposure on liking of that stimulus is faster, when the stimulus is simple [25] – resulting in a faster increase in liking with each repetition but also in a faster re-decline of liking of that stimulus. Thus, in the study a hand, very simple stimuli could have led to boredom, diminishing processing fluency and leading to more pronounced component activation after repetitive encounters.

While the results of the current study seem to replicate similar findings in the context of logo design [7], it is crucial to discuss ERP directionality of effects. During TWOI-1, Handy et al. [7] found increased P1 activation as a result of liked stimuli at central and parietal locations in comparison to disliked logos. Although the present study also revealed P1/N1 modulations, liked stimuli were associated with comparatively lower component activation and additional effects of perceived complexity seemed to play a crucial role (see Fig 1, panel a, affective ex-post approach). Interestingly, the volitional ex-post found a more pronounced N1 activation contrasting with previous findings [7].

In the time range of 200-300 ms (TWOI-2), Handy et al [7] presented evidence for less pronounced central N2 and a more pronounced parietal P2 components in response to liked logos in comparison to disliked logos. The evidence of the study at hand converges (affective ex-post approach) with previous findings as a less pronounced central N2 was found for liked stimuli (Fig 7, panel b) when stimulus material was perceived as complex. Additionally, we also found a more pronounced parietal P2 when stimulus material was complex. Contrasting evidence with previous studies was found for simple stimulus material. More specifically, the current analyses indicated a more pronounced N2 and a less pronounced P2 with increasing stimulus complexity. Comparing previous results [7] with the volitional ex-post approach, we find a similar less pronounced central N2 effect (see Fig 8, panel b). Furthermore, although parietal P2 modulations are also found in the volitional ex-post approach, the directionally was inversed (see Fig 8, panel b).

The diverging results of this study can be explained by the general inconclusive nature of liking effects directionality. For example, Righi et al. [12] also found similar effects to the study at hand comprising smaller P2 amplitudes in response to liked stimuli in go-stimuli in parietal locations – similar to stimulus material perceived as simple in this study. Additionally, as P2 (and N2) amplitudes have previously been shown to vary as a function of the intensity of negative and positive stimulus evaluation [14] this mechanism might also contributed to diverging result patterns.

Apart from potential differences in the intensity in participant reaction to different stimulus material, a methodological reason could be put forward to explain contrasting results. Arguably, some of the previously reported effects could be attributable to different stimulus material (e.g. different pictures) that were assigned to the groups of liked or disliked stimuli. E.g. in the study of Handy et al. [7] there is no apparent methodological control for stimulus material effects. This methodological issue is well-established in the field of psycho-linguistics and sometimes referred to as the “language as fixed-effect fallacy” which is sometimes generalised to “The Fixed - Effect Fallacy” [67]. Including the adequate error term (1|Prototype) is essential as stimuli represent a random sample from a broader population of potential stimuli. Failing to account for the respective random variability leads to inflated Type I error rates and overgeneralisation.

As a secondary aim, this study explored whether the application of specific design principles affects the neuro-physiological outcomes similar to beholder-based liking. Assessing manipulated prototype category effects showed higher liking values for the aesthetic category in comparison to the non-aesthetic category (see section *Results - Manipulation Check*). However, no effects of manipulated prototype category on ERP amplitudes or as in terms of a main effect or in interaction with ROI was found (see Table 5). Applying statistical control for the fixed-effects fallacy, seemed to be crucial applying the categorical ex-ante approach, as the same model without the (1|Prototype) error term shows prototype category effects (see supplementary material b) - a finding that was consciously omitted from the current article to avoid Error I inflation and overgeneralisation [67,68].

Interestingly, ERP Decoding revealed classification performance above chance levels for distinguishing manipulated aesthetic categories (see categorical ex-ante approach). This finding provides neural evidence that the aesthetic manipulation was successful, complementing the subjective increase in subjective liking. Importantly, decoding Haufe values correlated with ERP differences between non-aesthetic – aesthetic manipulated prototype category differences during both TWOIs. The above chance decoding, therefore, likely reflects systematic relative central-parietal amplitude differences associated with aesthetic processing.

Thus, the findings provide preliminary evidence for altered implicit neural processing of liked prototypes in response to the categorical prototype manipulation. The present finding seems to suggest that a small but significant subset of variables determining liking of NFB prototypes has been captured successfully by applying *Gestalt* principles – interpreting decoding accuracy the proportion of variance in liking accounted for by the applied design principles. Considering the decoding accuracy of 55.81 % it is obvious that a substantial proportion of variance remains unexplained. Thus, additional aesthetic principles and individual differences seem to play a crucial role in the design of aesthetic stimuli for NFB. This interpretation also aligns with the established principle of beholder-based nature of aesthetic experiences [e.g. 69].

All in all, the results of this study demonstrate that neural outcomes like ERP amplitudes are affected by design aesthetics, subjective liking and subjective complexity ratings. Respective effects can be considered a covariate in the context of NFB and they are understood only poorly at the moment. It remains to be explored by more specific future studies (see Fig 2) whether aesthetic experiences could improve NFB efficacy and efficiency directly e.g. via aesthetics effects on neural outcomes. Interestingly, indirect effects of aesthetic experiences on e.g. NFB time on task have already been demonstrated [6]. Moreover, user engagement has been shown before to be increased by aesthetic design [70] and deBeus and Kaiser [5] state increased engagement might mitigate the non-responder issue.

The current study confirms that beholder-based liking and complexity perceptions of NFB stimuli impacts phasic EEG responses. Thus, apart from the assumed NFB closed-loop effects of operant conditioning information presented on the screen, the way how the feedback information is presented affects the same outcome - i.e. brain activity patterns. Not only what is presented but how NFB information is transmitted matters.

Nevertheless, several limitations of this study are acknowledged that might have affected presented effects. First, ERP study material consisted of static images and did not include dynamic stimuli that would have more-closely resembled real-world NFB applications. While picture stimuli are the most common stimuli in the context of ERP studies [see e.g., 71], including video material and analysing ERP in response to e.g. categorical NFB changes (e.g. colour changes) could have improved external validity of this study. Secondly, as Gestalt principles seemed to explain only a small proportion of the variance of aesthetic experiences, other design principles should have been taken into consideration in the study at hand. E.g. fractal patterns have been shown to affect aesthetic experiences consistently [72], suggesting that the stimulus material in the present study could have been further optimised by application of corresponding design principles.

Taken together, the results speak in favour of the hypothesis of neurophysiological effects of beholder-based liking and complexity. The results further support the notion that reaction to NFB stimulus design is fast (100-200 ms after stimulus onset) and implicit (as liking and complexity rating were taken after study procedures were completed). Thus, indeed, feedback design might be an important covariate in the NFB closed-loop. Future studies should consider the character of design effects on further downstream variables.

## Acknowledgements

We would like to thank the Swiss National Science Foundation for funding the study at hand in the context of the project Advancing Neurofeedback in Tinnitus (ANT) Closing the loop with stimulus design and neural feature training registered under the project n° 208164.

## Conflict of interest

The authors declare that the research was conducted in the absence of any commercial or financial relationships that could be construed as a potential conflict of interest.

## Availability of data and materials

All relevant data and supplementary materials are available from: https://doi.org/10.5281/zenodo.18681583.

## Author Contributions

AN: Conceptualization, Project administration, Visualization, Writing – original draft, Writing – review & editing. DC: Resources, Writing – review & editing. PS: Methodology, Writing – review & editing. TK: Funding acquisition, Writing – review & editing. DRL: Resources, Writing – review & editing. PN: Writing – review & editing, Conceptualization. AS: Conceptualization, Project administration, Supervision, Writing – review & editing.

## Notes

### Competing Interest Statement

The authors have declared no competing interest.

